# Integrative analysis of the methylome and transcriptome of tomato fruit (*Solanum lycopersicum* L.) induced by postharvest handling

**DOI:** 10.1101/2023.10.17.562783

**Authors:** Jiaqi Zhou, Sitian Zhou, Bixuan Chen, Kamonwan Sangsoy, Kietsuda Luengwilai, Karin Albornoz, Diane M. Beckles

## Abstract

Tomato fruit ripening is triggered by the demethylation of key genes, which alters their transcriptional levels thereby initiating and propagating a cascade of physiological events. What is unknown, is how these processes are altered when fruit are ripened using postharvest practices to extend shelf-life, as these practices often reduce fruit quality. To address this, postharvest handling-induced changes in the fruit DNA methylome and transcriptome, and how they correlated with ripening speed, and ripening indicators such as ethylene, ABA, and carotenoids, were assessed. This study comprehensively connected changes in physiological events with dynamic molecular changes. Ripening fruit that reached ‘Turning’ (T) after storage under dark at 20°C, 12.5°C, or 5°C chilling (followed by 20°C rewarming), were compared to fresh-harvest fruit ‘FHT’. Fruit stored at 12.5°C, had the biggest epigenetic marks and alterations in gene expression, exceeding changes induced by postharvest chilling. Fruit physiological and chronological age were uncoupled at 12.5°C, as the time-to-ripening was longest. Fruit ripening at 12.5°C was not climacteric; there was no respiratory or ethylene burst, rather, fruit were high in ABA. Clear differentiation between postharvest-ripened and ‘FHT’ was evident in the methylome and transcriptome. Higher expression of photosynthetic genes and chlorophyll levels in ‘FHT’ fruit, pointing to light as influencing the molecular changes in fruit ripening. Finally, correlative analyses of the -omics data putatively identified genes regulated by DNA methylation. Collectively these data improve our interpretation of how tomato fruit ripening patterns are altered by postharvest practices, and long-term are expected to help improve fruit quality.

## Introduction

Postharvest handling approaches are commonly used to extend tomato fruit shelf-life; however, they often negatively affect fruit quality attributes. Although a longer shelf-life reduces the rate of fruit deterioration and postharvest losses while making produce suitable for commercial sale, the unintended negative effects on quality can lead to rejection by the consumers, creating postharvest waste ^1^. Understanding the mechanisms of postharvest-induced changes in tomato through fruit physiology and molecular biology is a first step towards finding a solution for postharvest loss and waste ^2^.

Harvesting tomato fruit before full-ripening is an efficient approach to extend their shelf-life. However, the loss of the energy and nutrient support from the mother plant often causes off-the-vine fruit to be suboptimal in quality, negatively influencing fruit sugar-to-acid ratio, volatile profiles, texture, and weight ^3–6^. Depending on the postharvest storage conditions, i.e., temperature, light, dark, humidity, CO_2_, and O_2_ concentration, fruit ripening and the development of quality traits are differentially affected ^7^. Conversely, fruit ripened on the vine can import sugars and other compounds for an extended period and are exposed to a longer period of sunlight which is important to fruit quality ^8^.

Cold storage is used to slow down senescence and preserve quality in harvested fruit by reducing respiration, biochemical reactions, the rate of fungal infestation and water loss ^2^. However, tomato and other tropical and subtropical crops are sensitive to low temperatures. Postharvest chilling injury (PCI) widely occurs when sensitive produce are stored at temperatures below 12.5°C ^9–11^.

Tomato fruit stored below 12.5°C may show symptoms of PCI upon rewarming to room temperature such as abnormal firmness and texture, uneven ripening, fruit surface pitting, and spoilage from fungi ^12^. The severity of PCI symptoms depends on the time-temperature combination and preharvest factors ^13^.

The current understanding of the molecular basis of fruit development, ripening, and senescence is highly developed in tomato, even if there remain many unanswered questions. The regulation of fruit ripening mechanisms focuses on hormones, mainly ethylene, but also in recent years, ABA, jasmonic acid, cytokinin, gibberellins, and auxin. The rapid increase in ethylene is a well-established and critical feature of climacteric fruit ripening ^14–16^, but ABA has recently been discovered as a positive regulator of tomato fruit ripening ^17,18^. ABA is also a stress hormone, responsive to abiotic environments like temperature and water conditions. The mechanism of hormone interplay in fruit is still unclear. The current hypothesis is that ABA works prior to ethylene and activates tomato fruit ripening ^19^, but the relationship between the role of ABA and tomato fruit ripening after harvest is unknown.

A critical role for DNA demethylation in governing tomato fruit ripening and hence quality, has also been recognized. Demethylation events occur at the promoter regions of ripening genes, presumably controlling transcription factor binding, thereby dictating if genes will be turned on/off^20^. Active DNA demethylation is enacted by DNA glycosylases, of which SlDML2 is the most important in tomato, as silencing *SlDML2* halts ripening ^21^. Chilling stress inhibits *SlDML2* expression, suppressing ripening-associated demethylation; however, this action is partially reversed when fruit are rewarmed ^22^. Changes in tomato fruit DNA methylation levels due to chilling correlate with fruit flavor loss and variation in the transcriptional levels of key ripening genes ^23^. Other epigenetic modifications also affect DNA demethylation ^24^, and this epigenome remodeling can collectively change fruit shelf life and quality ^5,25^.

The widespread reprogramming that occurs during ripening can be explored using -omics scale research, where multiple biological pathways can be simultaneously explored to systematically unravel the underlying mechanisms ^26^. Transcriptomic analysis has enabled an understanding of key ripening pathways under varied postharvest conditions ^25^. DNA methylomics analysis can precisely pinpoint changes in methylation status at loci under certain conditions. Individually, - omics studies like transcriptomics and methylomics can be used to explore global differences, and generate co-expression networks with key markers highlighted across treatments ^27^. Integrating these data can find correlations among epigenetic and transcriptional changes, pointing out potential regulatory mechanisms of key biological process ^28^.

In this work, we studied how postharvest handling, i.e., off-the-vine ripening and low temperature storage affect tomato ripening and quality, through accessing the fruit transcriptome and methylome, and by studying ripening hormones and quality traits. Comparisons were made on fruit at the same developmental stage but that underwent different postharvest storage simulating conditions used in industry. Integrative analysis was used to connect fruit ripening physiology and events at the epigenomic and transcriptomic level. Long term, postharvest biomarkers for future targets could be explored.

## Material and Methods

### 2.1 Plant growth

*Solanum lycopersicum* L. cv. ‘Micro-Tom’, an experimental model cultivar for postharvest studies was used. ‘Micro-Tom’ seeds were from the Tomato Genetics Research Center at UC Davis. Germination and plant growth methods were as described previously^5^. Postharvest treatments were done on fruit randomly harvested from over one hundred plants in 2020, 2021, and 2022.

### 2.2 Fruit sampling and postharvest treatments

Fruit were sampled at two developmental stages - Mature Green (MG) and Turning (T), as described by Zhou *et al*., (2021) ^5^ (Fig. S1). Harvested fruit were washed with 0.27% (v/v) sodium hypochlorite for 3 min and air dried. Fruit harvested at MG (named as ‘FHM’) were stored in the dark and analyzed when they reached Turning ‘T’ after storage at (1) 20℃ (named as ‘20T’); (2) 12.5℃ (named as ‘12.5T’), and (3) 5℃ for two weeks followed by rewarming at 20℃ (named as ‘5T’). The control group is the fresh harvested Turning fruit (‘FHT’). MG fruit were also analyzed after storage at 5℃ for two weeks (‘5M’). Three biological replicates, each consisting of a pool of six randomly selected fruit pericarps, were sampled for whole-genome bisulfite sequencing, RNASeq, carotenoids and ABA assays.

### 2.3 Whole genome bisulfite sequencing (WGBS)

#### Genomic DNA extraction

Genomic DNA was isolated using the Qiagen® DNeasy Plant Mini Kit. Due to the high carbohydrates of ripening tomato fruit, the procedures were modified according to the manufacturer’s protocol to increase DNA yields and quality. The extraction for each sample was started with a duplicate sample material, and one extraction of 100 mg frozen fresh fruit powder were added into the buffer AP1 and P3 followed by QIAshredder columns, respectively. The flow-through from the duplicate extractions was pooled together, and after adding AW1, all mixtures were loaded into one DNeasy Mini spin column. In the final elution, the AE buffer was preheated at 65℃ and incubated for 30 min for the best elution efficiency. The isolated DNA was further purified using the DNA Clean & Concentrator-5 (Zymo Research Corp., Irvine, CA, USA). The quality of DNA was assessed on the 0.8% (w/v) agarose gel, a microvolume spectrophotometer and a Bioanalyzer (Agilent, Santa Clara, CA, USA).

#### Methyl-Seq Library preparation and sequencing

The bisulfite conversion of sonicated genomic DNA fragments was carried out based on the instructions provided in the EZ DNA-methylation lightning Kit (Zymo Research Corp., Irvine, CA, USA). The libraries were made using the Accel-NGS Methyl-Seq DNA library kit (SWIFT Biosciences, Ann Arbor, MI) and quality checked using the Bioanalyzer. The libraries were sequenced using the NovaSeq PE 150 at the UC Davis Genome Center DNA Technologies & Expression Analysis Core.

#### Data processing

The sequencing reads were first quality checked on FastQC ^29^, and all libraries passed quality control requirements, after adaptor trimming using Trimmomactic ^30^. The bisulfite conversion rates were calculated by aligning reads to the unmethylation chloroplast genome, and the conversion rates for all libraries were more than 97% ^31^. The trimmed reads were aligned to the tomato genome assembly SL 3.0 (Sol Genomic Network) using Bismark ^32^. The multi-aligned reads were deduplicated to remove PCR bias. Methylation extraction was conducted to calculate the methylated status of each sequenced cytosine and extracted by CpG, CHH, and CHG contexts respectively. The visualization of the DNA methylation status and correlation between each library were performed in SeqMonk (https://www.bioinformatics.babraham.ac.uk/projects/seqmonk/). The final Bismark output text files were imported to R (R Core Team, 2020). The differentially methylated regions (DMRs) and differentially methylated genes (*p* < 0.05) were extracted using MethylKit ^33^ and were annotated using the Genomation package ^34^. The DMRs were defined by a threshold of *p* < 0.05, the difference of the methylation percentage > 10, using a 200-bp sliding window.

### 2.4 RNASeq library preparations and sequencing

#### RNA isolation

Fruit pericarp were frozen by liquid nitrogen and stored at −70°C upon sampling. Total RNA was isolated from around 100 mg fruit powder using a Trizol-based protocol. RNA quality and integrity were assessed by microvolume spectrophotometer and 0.8% (w/v) agarose gel electrophoresis. The mRNA was isolated from total RNA using NEBNext® Poly(A) mRNA Magnetic Isolation Module.

#### 3’ DGE RNASeq library construction and sequencing

The libraries were built using Strand-specific mRNA-library prep kits (Amaryllis Nucleics, Oakland, CA). All libraries that passed the quality check conducted by Novogene, were pooled into one lane, and sequenced by HiSeq PE150. The raw sequencing reads were trimmed for removing adaptors using Trimmomatic ^30^, and quality checked by FastQC ^29^. The reads alignment was processed by STAR ^35^ based on the tomato reference genome SL4.0 (Sol Genomic Network). Visualization of the aligned reads were performed in SeqMonk. The aligned reads were imported to R and processed by the package FeatureCounts ^36^ to obtain the read count of each gene. Data normalization and clustering were performed before extracting differentially expressed genes (DEGs) by EdgeR ^37^. The input of the GO terms was downloaded using the BioMart tool at Ensembl Plants (http://plants.ensembl.org/biomart/martview/) for both DEGs and DMGs annotation. The functional enrichment analyses including Gene ontology by GOseq ^38^, and KEGG by Gage ^39^ were conducted.

### 2.5 Bioinformatics analysis

#### Co-expression network

Gene modules were identified using the WGCNA under the R environment ^40^, from 15 samples (‘5T’, ‘12.5T’, ‘20T’, ‘FHT’, and ‘5M’, each with three biological replicates) in the RNASeq data. The correlation network analysis includes 2255 significant genes identified in at least one comparison between postharvest Turning fruit and ‘FHT’, i.e., ‘5T’ vs. ‘FHT’, ‘12.5T’ vs. ‘FHT’, and ‘20T’ vs. ‘FHT’. The power (soft threshold) was determined by the pickSoftThreshold function in the WGCNA package. An unsigned network was constructed using automatic network construction, with minModuleSize of 30 and mergeCutHeight of 0.25. The eigengene expressions were obtained, and Pearson’s correlation coefficient (PCC) represented by *r* was used to calculate the correlation between each module and treatment group. Furthermore, the top 1000 strongest connections, identified as gene pairs with the highest edge weight, were further imported to Cytoscape (version 3.9.1) ^41^ for network visualization.

#### Hub genes

Hub genes in each module were identified through a multi-criteria approach. First, genes with the top 10% intramodular connectivity were selected. The intramodular connectivity was calculated using the function intramodularConnectivity in WGCNA. Second, the selected genes were further filtered for the absolute geneModuleMembership (KME) value greater than 0.9, where the KME value was calculated by signedKME in WGCNA package. The filtered genes were then combined with the top 1000 strongest connections identified in section above to find those that overlapped. The overlapped genes were identified as the hub genes that are strongly associated with and highly connected within candidate modules.

#### Gene ontology visualization using GoFigure

Go Figure ^42^, a Python package, was used for GO visualization. The GO categories and the associated overrepresented *p*-values for each module were imported into the program to create the plots.

#### Correlations between gene associated DNA methylation regions and gene expression levels

The correlation between gene expression and DNA methylation levels was calculated for each differentially methylated regions (DMRs) determined in three methylation contexts, i.e., CG, CHG and CHH. For each DMR, the RNASeq data with three biological replicates were used as the gene expression levels, and the average DNA methylation percentage across all contexts was used as the DNA methylation levels. Correlations were calculated separately if there were multiple DNA methylation sliding windows identified for one gene. The PCC represented by *r* and its *p*-values were calculated to indicate the strength of correlation.

For genes in specific pathways, the correlation between their gene expression and DNA methylation levels was examined. The DNA methylation levels were based on the regions surrounding the gene, including the 3 kb upstream and gene coding regions. The correlation was indicated by *r*, and statistical test indicated by *p*-values were summarized in tables.

### 2.6 Fruit carotenoids

Carotenoids extractions and assay were done as previously described ^43^ with some modifications. Frozen tomato tissue (0.2-0.4 g) was extracted with 20 mL HEA (2:1:1 hexane: ethanol: acetone, v/v/v) containing 0.1% (w/v) butylhydroxytoluene. The extracted carotenoids were covered with aluminum foil to avoid light exposure. The extraction was repeated to collect all supernatants after centrifugation until the tomato tissue is colorless. The homogenized extract was incubated for 15 min in the dark at room temperature, and 15 mL distilled water was added, and the extract was incubated further for 15 min. The organic phase was separated and evaporated under high pressure N_2_ until dry. Carotenoids contents were analyzed using high performance liquid chromatography (HPLC; Agilent 1100, Hewlett-Packard-Strasse, Germany). The dried extract was dissolved in 1-mL of the mobile phase (10: 5: 85 dichloromethane: acetonitrile: ethanol, v/v/v) ^44^ and filtered through a 0.22 μm nylon membrane. The sample (20 μL) was injected into the HPLC equipped with a YMC-C30 reversed-phase column (25 mm × 4.6 mm,5 μm, YMC Co., Kyoto, Japan). The flow rate was 1 mL/min at ambient temperature (25°C), and the absorption of each compound was detected with a UV–Vis detector. Absorption spectra for the main peaks were 285 nm for phytofluene and 450 nm for lycopene, β-carotene, and lutein. A chromatographic run lasted 65 min. Each carotenoid was identified by the retention time compared with the external standard. Phytofluene standards was purchased from CaroteNature GmbH, (Lupsingen, Switzerland). Lycopene (9879), β-carotene (22040) and lutein (07168) standards were purchased from Sigma-Aldrich, USA.

### 2.7 Fruit abscisic acid (ABA) extraction and ELISA-antibody kit analysis

The ABA extraction methods were modified from a previous study ^45^. Approximately 50 ∼ 100 mg of frozen tomato tissues were ground in liquid nitrogen and used for the extraction. One milliliter of the extraction buffer (80% (v/v) methanol (methanol: water: acetic acid (80:19:1, v/v) with 100 mg/L butylated hydroxytoluene (BHT)) was added in each sample and the incubation was conducted at 4℃ in the dark. After 24 hours, the supernatant and pellet were separated by centrifuging, and the incubation was repeated using another 1 mL extraction buffer for an additional hour. All supernatants were collected and dried in a speed vac. The dry pellet was dissolved in 99% (v/v) methanol (Methanol: water: acetic acid (99:1, v/v) with 100 mg/L BHT. The dissolved pellet was added with 900 μL 1% (v/v) acetic acid, loading into the Sep-pak C18 reverse phase columns (Waters, USA). The column was washed with 3 mL of 20% (v/v) methanol following elution by 3 mL of 80% (v/v) methanol (methanol: water: acetic acid (80:19:1, v/v) with 100 mg/L BHT. The eluted samples were dried, and the pellet was dissolved with 50 μL methanol and 450 μL Tris-buffered saline (TBS) buffer. The extracts were diluted 20-fold using TBS buffer before the Phytodetek® ELISA-plant ABA kit assay (Agdia, Inc., Elkhart, IN).

### 2.8 Fruit difference of absorbance (DA) index and color assay

A DA meter® (TR Turoni, Italy) was used for the non-destructive assessment of fruit chlorophyll content, while a colorimeter (Konica Minolta, Tokyo, Japan) was used for measuring objective color. The color was used as the determinant for fruit developmental stage in this study. Each fruit was assessed twice at the equatorial regions of the skin according to Albornoz *et al*., (2019) ^13^. At least twenty tomato fruit were measured in each treatment group.

### 2.9 Fruit postharvest gas analysis-ethylene and respiration rates

Tomato fruit at the mature green stage were harvested in the morning and stored under different temperatures. The gas assays were performed daily at a similar time. Around one hundred grams of fruit were pooled in one jar as one biological replicate. Six biological replicates, each with at least two repeated assays (technical replicates), were included. The fruit were placed in a sealed 450 mL glass jar for 30 to 60 min each day, and gas was extracted for assaying ethylene and CO_2_. Ethylene was measured by a gas chromatograph, and carbon dioxide was assayed by a CO_2_ analyzer ^13^.

### 2.10 Validation of the RNASeq identified DEGs using qRT-PCR

Fruit harvesting and postharvest treatments were repeated to consider pre-harvest environmental factors affecting the fruit transcriptome. Tomato plants were grown in the greenhouse at UC Davis, CA in 2023. Postharvest treatments were performed on fruit randomly harvested over 50 plants. Six fruits were randomly selected and pooled together to form one biological replicate. Three biological replicates and four technical replicates were included. Fruit pericarp samples were frozen into liquid nitrogen and stored at −70°C upon sampling. Total RNA was isolated from 100 mg fruit powder using a Trizol-based protocol. RNA quality and integrity were assessed by microvolume spectrophotometer and 0.8% (w/v) agarose gel electrophoresis. cDNA libraries were reverse transcribed, and qRT-PCR were performed according to our previous study ^5^. The *SlFRG27* (*Solyc06g007510*) was the internal control reference gene for all tested genes ^46^. The ‘FHT’ was used as the control to compare with each postharvest treatment.

## 3. Results

### 3.1 Postharvest ripening fruit quality and methylome

Fruit were harvested at mature green (MG) and allowed to ripen at 20°C, 12.5°C, 5°C and 5°C plus rewarming at 20°C as described previously ^5^. There were two MG groups, i.e., fruit fresh-harvested at MG (‘FHM’), and ‘FHM’ stored at 5℃ for two weeks (‘5M’). There were four Turning fruit groups: three were ripened postharvest, i.e., fruit were harvested at ‘FHM’ and then stored at 20°C (‘20T’), 12.5°C (‘12.5T’), and 5°C plus rewarming at 20°C (‘5T’), and the fourth group is fresh harvested Turning (‘FHT’) that ripened on-the-vine (Fig. S1).

The quality traits in the fruit samples, were assessed in our previous work ^5^ and included fruit color, reducing sugars, total soluble solids, starch, titratable acids, and firmness. These quality data were presented in a principal component analysis (PCA) plot (Fig. 1A). Fruit quality was mainly clustered by ripening stages (color), and among the four Turning groups, ‘5T’ was the most distinct and presumably the worst quality profile ^5^ from others.

**Figure 1.**
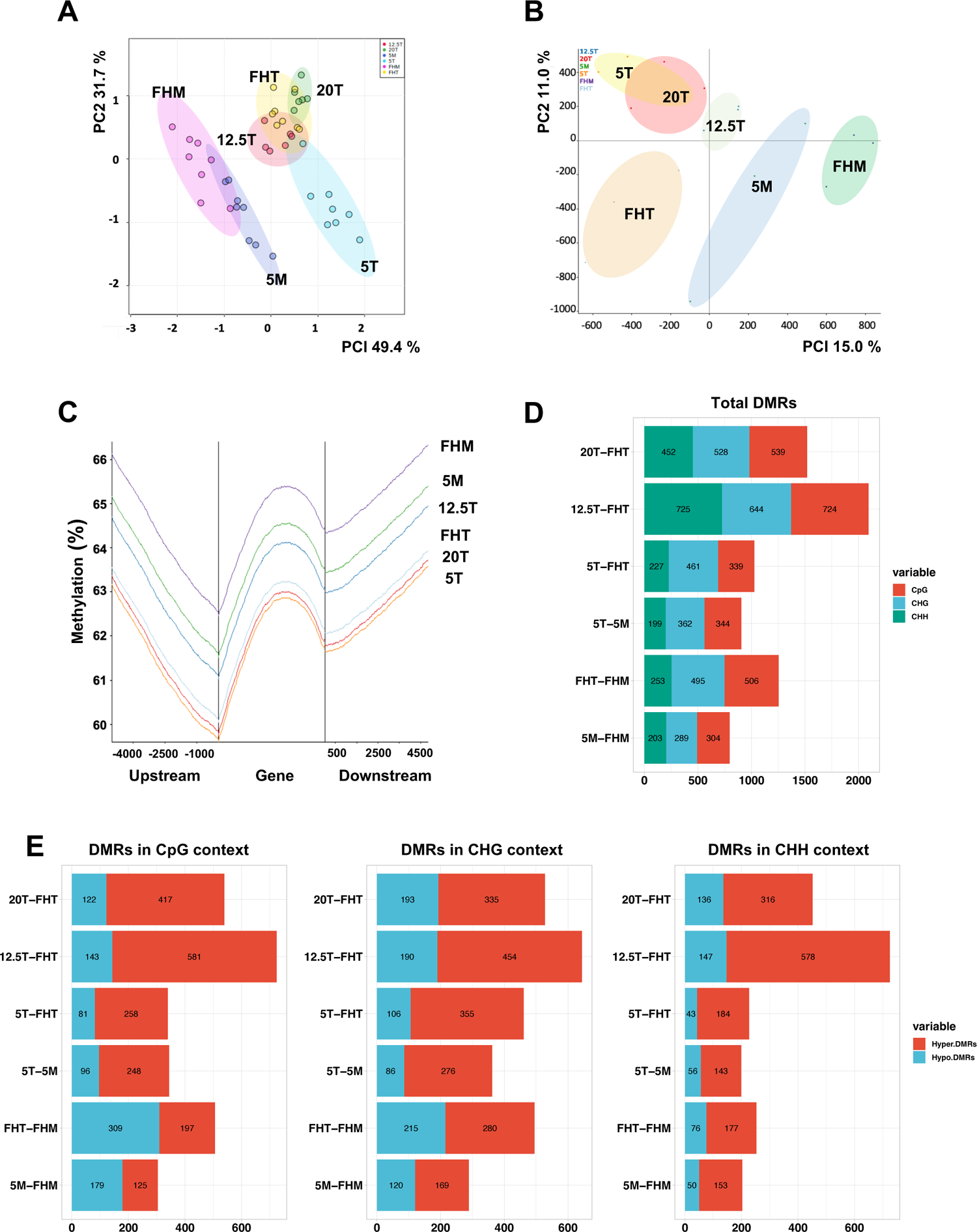
Analysis of the postharvest tomato fruit quality and methylome. **(A)** Principal Component Analysis (PCA) of the fruit quality parameters. (**B**) PCA of the fruit methylome. **(B)** Methylation in the genic regions shown as a change in methylation percentage using a 200C (Cytosine) sliding window. **(D)** Proportion of differentially methylated regions (DMRs) in each comparison at CG, CHG and CHH contexts. **(E)** Among the number of DMRs in **(D)**, there are hypo- and hyper-DMRs when compared to ‘FHT’.

To determine the influence of various postharvest treatments on fruit methylation, context specific methylation levels for each treatment were assessed (Figs. S2-S3, Table S1). The methylome of green fruit (‘FHM’ and ‘5M’) was similar to each other and distinct from Turning fruit (Fig. 1B).

Within the Turning fruit, those ripened postharvest, i.e., ‘20T’, ‘12.5T’ and ‘5T’, clustered away from ‘FHT’, suggesting that ripening after harvest, regardless of storage temperature, affected the fruit methylome.

When comparing the quality and DNA methylation PCA (Figs. 1A-B), incongruity was seen between ‘FHT’ and ‘5T’. ‘FHT’ has similar quality traits as the off-the-vine ripening ‘20T’ but a different methylation profile, whereas ‘5T’ has a similar methylation status to ‘20T’ but distinctly lower quality. We anticipated greater methylation marks on genes in cold-stored fruit, and the ‘12.5T’ would be more similar to other Turning fruit, but in contrast, our data shown ‘12.5T’ was even more similar to ‘5M’, comparing data derived from the whole genome (Fig. 1C). The relative DNA methylation levels within genes were: ‘FHM’ > ‘5M’ > ‘12.5T’ > ‘FHT’ > ‘20T’ > ‘5T’ (Fig. 1C).

To understand how the DNA methylation status varied between two samples, pair-wise comparisons were performed, and the differential methylated regions (DMRs) were identified. The comparisons were as follows: (1) ‘5M’ vs. ‘FHM’ for chilling effect, and (2) ‘5T’ vs. ‘5M’ for the influence due to ripening and rewarming. By comparing each postharvest ripened fruit to the ‘FHT’, i.e., (3) ‘5T’ (4) ‘12.5T’ and (5) ‘20T’, DMRs due to off-the-vine ripening at the respective temperatures could be inferred (Figs. 1D-E). The total DMRs showed that the ‘12.5T’, was the most unusual. The ‘12.5T’ had the highest number of DMRs compared to ‘FHT’ in all methylation contexts (Fig. 1D). Further, most of the DMRs in ‘12.5T’ were hypermethylated compared to ‘FHT’ (Fig. 1E) and were in non-gene associated regions (Fig. S4).

Taken together, these data suggest that (1) the different postharvest ripening processes significantly altered the fruit DNA methylome, and (2) the low but non-chilling temperature storage (‘12.5T’) led to great changes at the methylome, presumably through its unique effect on fruit development and ripening compared to other storage temperatures.

### 3.2 Functional analysis of the differential methylated regions (DMRs)

In this work, differentially methylated genes (DMGs) were defined as having DMRs around the gene body or 3 kb upstream promoter regions ^47^. With the fruit transcriptomic data, DMGs expressed in the experimental fruit were filtered and summarized in Table S2. The Venn plot in Fig. 2A shows the overlapping or unique DMGs among postharvest Turning fruit compared to ‘FHT’. The ‘12.5T’ treatment had the greatest number of DMGs compared to the ‘FHT’, i.e., 91 unique genes, 37 overlapped with ‘20T’, 31 were shared with ‘5T’, and 47 overlapped with both ‘20T’ and ‘5T’.

**Figure 2.**
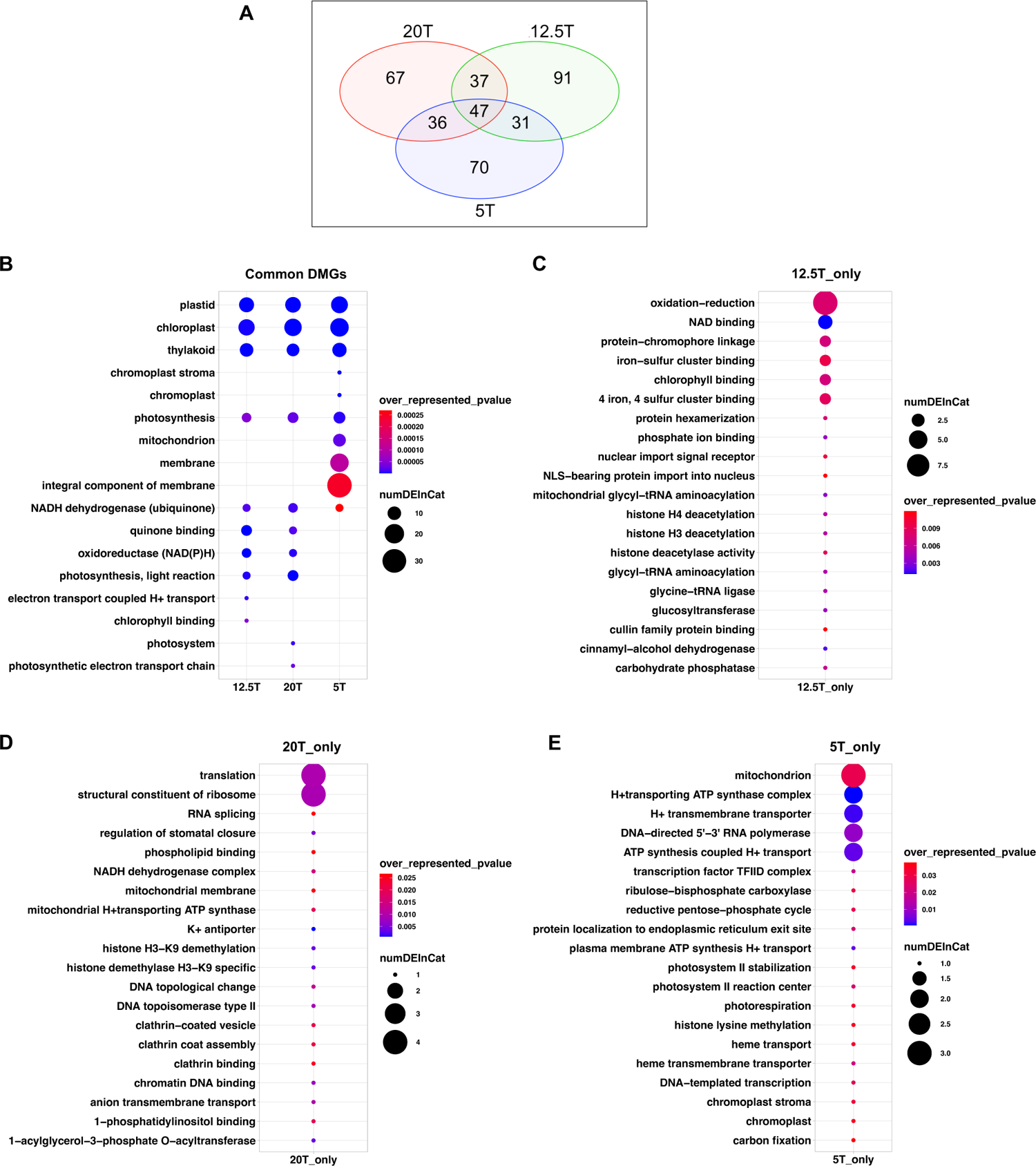
Differential methylated genes (DMGs) and Gene Ontology term analysis of the DMGs. **(A)** Venn plot of the DMGs using ‘FHT’ as the control. The number of different and overlapped DMGs were shown in the center of the plot. **(B)** GO analysis for the DMGs in **(A)**, and the top GO terms were shown in the plot. The unique GO terms in ‘12.5T’ were in **(C)**, in ‘20T’ were in **(D)** and in ‘5T’ were in **(E)**.

The enriched Gene ontology (GO) terms of all three treatments are plastids (both ‘chloroplast’ and ‘chromoplast’), ‘photosynthesis’ and ‘NADH dehydrogenase’ (Fig. 2B). This implies that genes expressed during the fruit chloroplast to chromoplast transition may be modulated by DNA methylation. Figs. 2C-E show the GO terms associated with each postharvest treatment. The unique GO terms for ‘12.5T’ were enriched in ‘oxidation-reduction’, ‘NAD binding’, ‘histone deacetylation’ and others. The ‘20T’ were related to ‘translation’ and ‘histone demethylase activity’, and the ‘5T’ was enriched with the ‘mitochondrion’ and ‘ATP synthase’ (Details of GO terms are in Table S3).

### 3.3 Postharvest ripening fruit transcriptome

Variation in gene methylation may have consequences for gene expression and downstream physiological processes. To examine this, we profiled changes in tomato fruit transcriptome and quality parameter assessments. RNASeq analysis indicated that 16,129 genes were expressed in fruit after filtering out the lowly-expressed genes. We focused on the fruit ripened postharvest and compared them to fruit ripened on the vine (‘FHT’). Postharvest ripened fruit were more like each other and differed from ‘FHT’ (Figs. 3A and 3B). Although the fruit ripened after cold storage, i.e., ‘5T’, had quality biomarkers that differed from ‘20T’ (Fig. 1A), when comparing mRNAs, these fruit were more similar. Therefore, the most recent temperature exposure influenced the fruit transcriptome, erasing the effects of the prior chilling event ^5,22^.

**Figure 3.**
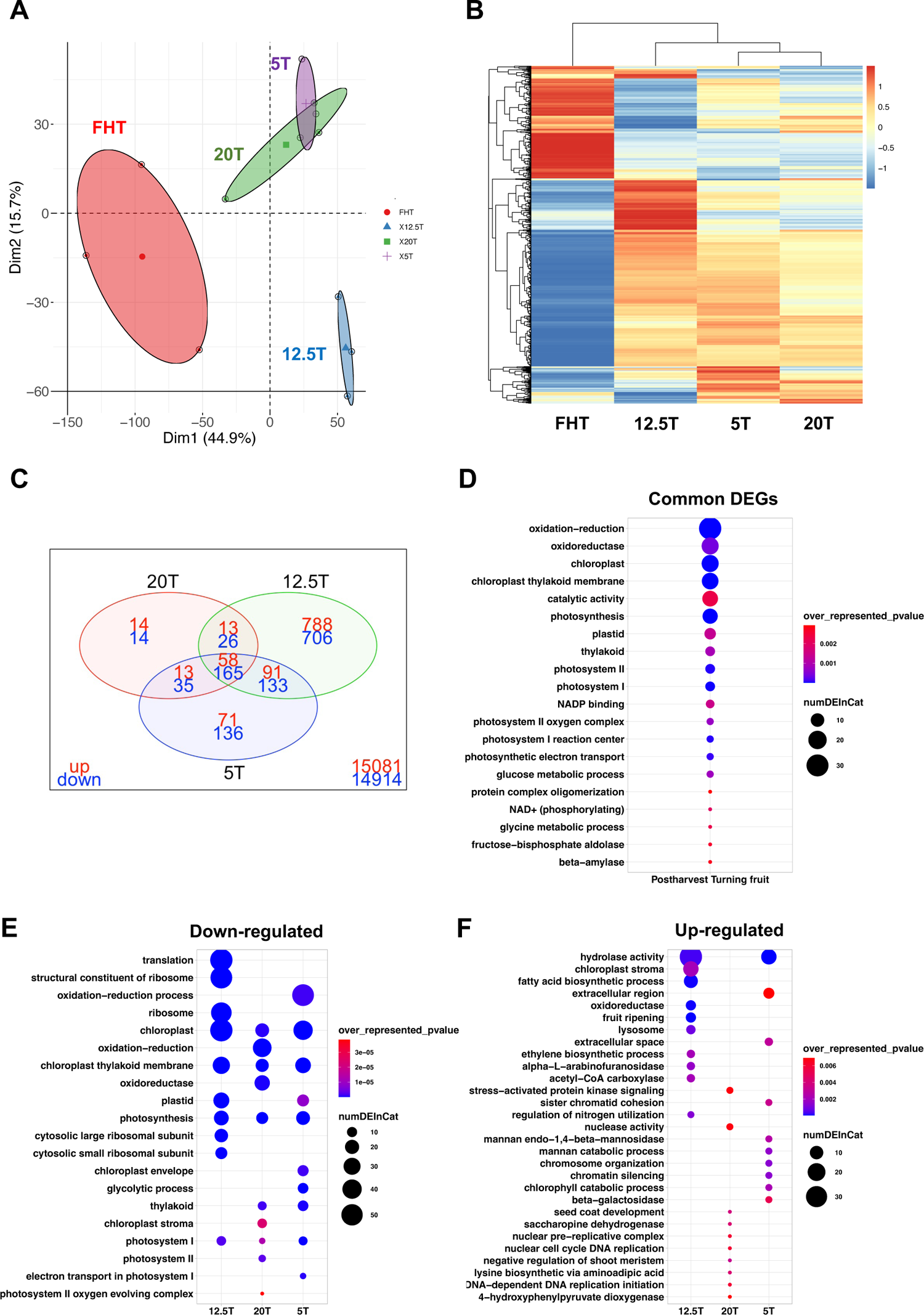
Tomato fruit transcriptome. (**A**) PCA of the transcriptome in ‘Turning’ fruit, i.e., ‘FHT’, ‘20T’, ‘12.5T’ and ‘5T’. (**B**) Hierarchical clustering and heatmap were drawn for the most variable 1000 genes based on their expression levels. Genes in red and blue represent highly- and lowly-expressed genes, respectively. (**C**) Venn plot of the differentially expressed genes (DEGs) in pair-wise comparisons. The numbers in red and blue represent upregulated and downregulated DEGs compared to ‘FHT’, respectively. (**D**) Gene Ontology enrichment analysis of common DEGs (postharvest fruit compared to ‘FHT’.) The top 20 GO terms (*p* < 0.05) were shown. (**E**) **(F)** when the postharvest Turning, i.e., ‘12.5T’, ‘20T’ or ‘5T’ was compared to the ‘FHT’, the top representative GO terms (*p* < 0.05) for downregulated genes were shown in **(E)** and upregulated genes in **(F)**.

The differentially expressed genes (DEGs) in the pair-wise comparisons were identified (Table S4). The ‘12.5T’ had the highest number of DEGs (1030 up and 950 down) compared to all other groups (Fig. 3C.), indicating that the low but non-chilling temperature strongly regulated gene expression during fruit ripening. This is surprisingly consistent with the DNA methylation data for these fruit. The ‘20T’ fruit was similar to ‘FHT’, having the lowest number of DEGs, most likely related to early harvest and dark storage treatments.

Functional enrichment analysis of the common DEGs (58 up- and 165 down-regulated) for all postharvest turning fruit (Fig. 3C), vs. ‘FHT’ was summarized (Table S5), and the top GO terms are shown in Fig. 3D and the KEGG analysis was in Fig. S14. Terms related to photosynthetic and respiration-related activities were prevalent. This finding aligns with the functional analysis of the DMGs which collectively indicate physiological alterations in energy capture and use in postharvest ripened fruit compared to vine-ripened fruit (Fig. 2B). These data also illustrate potential correlations between DNA methylation and gene expression, and their effect on biochemical mechanisms. In addition, the gene associated with ‘beta-amylase’ activity were enriched (Fig. 3C), specifically *beta-amylase 8* is differentially expressed among Turning groups. Higher *beta-amylase 8* expression in all postharvest fruit compared to ‘FHT’, was also validated by qRT-PCR (Fig. S15), indicating that starch degradation may be more active during the off-the-vine ripening process, which corresponds to the reduced starch in the postharvest fruit ^5^.

Enriched GO terms specific to each postharvest group were summarized in Table S5. When pooling the down- or up-regulated GO terms specific to each group, the shared or unique terms were explored (Figs. 3E and F). Because of the large number of DEGs in the ‘12.5T’ fruit, substantial differences were seen and the DEGs were enriched for ‘translation’ (Fig. 3E) and ‘fruit ripening’ (Fig. 3F). Changes at the post-translation levels, in addition to methylation variations may be prominent in ‘12.5T’ relative to ‘FHT’. While ‘12.5T’ fruit had the same objective color and similar ripening characteristics to other Turning fruit ^5,^ this was not reflected at the transcriptional level. It may be important to explore differences in fruit chronological age (the time fruit took to reach the certain ripening stage) and relative physiological age ^48^ (‘Turning’ stage in this work), under varied environmental conditions.

Many photosynthesis-associated pathways were downregulated in fruit ripened postharvest compared to ‘FHT’ fruit (Fig. 3E). If these genes regulate fruit photosynthesis, then this biochemical process may still be active in the on-the-vine growing tomato fruit, but not in harvested fruit stored in the dark. Among the upregulated genes that differentiated postharvest fruit from ‘FHT’, ‘hydrolase activity’ is enriched in both cold stored fruit, i.e., ‘12.5T’ and ‘5T’ (Fig. 3F). In addition, the upregulated GO enrichment has no GO terms shared across three treatments, unlike that were downregulated.

### 3.4 Gene expression co-expression network by WGCNA

We used weighted gene co-expression analysis (WGCNA) to identify gene modules related to specific postharvest storage conditions. The DEGs from the comparisons of postharvest Turning (i.e., ‘20T’, ‘12.5T’, ‘5T’) to the fresh-harvested Turning (‘FHT’) were pooled together. The 2255 unique genes as the input dataset, were clustered as six module eigengenes (ME; Fig. S5A), i.e., turquoise (993 genes), blue (539), brown (358), yellow (182), grey (128) and green (55).

ME_turquoise and ME_blue are distinct with only limited overlapping genes (Fig. S5A). Relationship between each gene module and the fruit treatments were identified (Fig. S5B). Heatmaps for these module-trait relationship were developed, and their correlation (*r*) and significance (*P-*value) were indicated in each block. The genes in the ME_turquoise only correlated with ‘FHT’ (*r* =0.82, *p* < 0.001), and not for postharvest-ripened fruit. Genes in the ME_blue were positively correlated to ‘12.5T’ (*r* =0.78, *p* < 0.001). Integrating these observations with that of transcriptome PCA, indicate that ‘FHT’ and ‘12.5T’ have distinct gene expression patterns even though they are in the same ‘color-indicated’ ripening stage.

The genes in each ME were annotated using GO terms ^42^ (Fig. S6). ME_turquoise was enriched in genes for ‘respiration’, ‘photosynthesis’, and ‘translation-related’ activities. ME_blue was enriched in genes encoding secondary metabolites, while ME_brown was enriched in photosynthesis and starch biosynthesis genes.

The gene network of each module is presented in Fig. S5. The clustering of genes helps to identify the hub genes with high connectivity to other genes (Table S6). The hub genes potentially work upstream in fruit transcriptome response to postharvest treatments, so they could be the candidate targets to study postharvest fruit ripening biology ^49^.

### 3.5 Fruit carotenoids and abscisic acid (ABA) content

To connect the molecular and biochemical events with quality parameters, fruit physiological processes related to ripening were assayed. Fruit carotenoids, including lycopene, β-carotene, lutein and phytofluene were assessed in Turning fruit. The ‘12.5T’ had relatively high carotenoids, and uniquely its β-carotene was 2.6-fold higher than ‘FHT’ (Fig. 4A). There was high within-group variability in the carotenoids data, indicating that postharvest treatments interacting with preharvest factors, altered fruit quality changeably, while influencing fruit quality ^50^.

**Figure 4.**
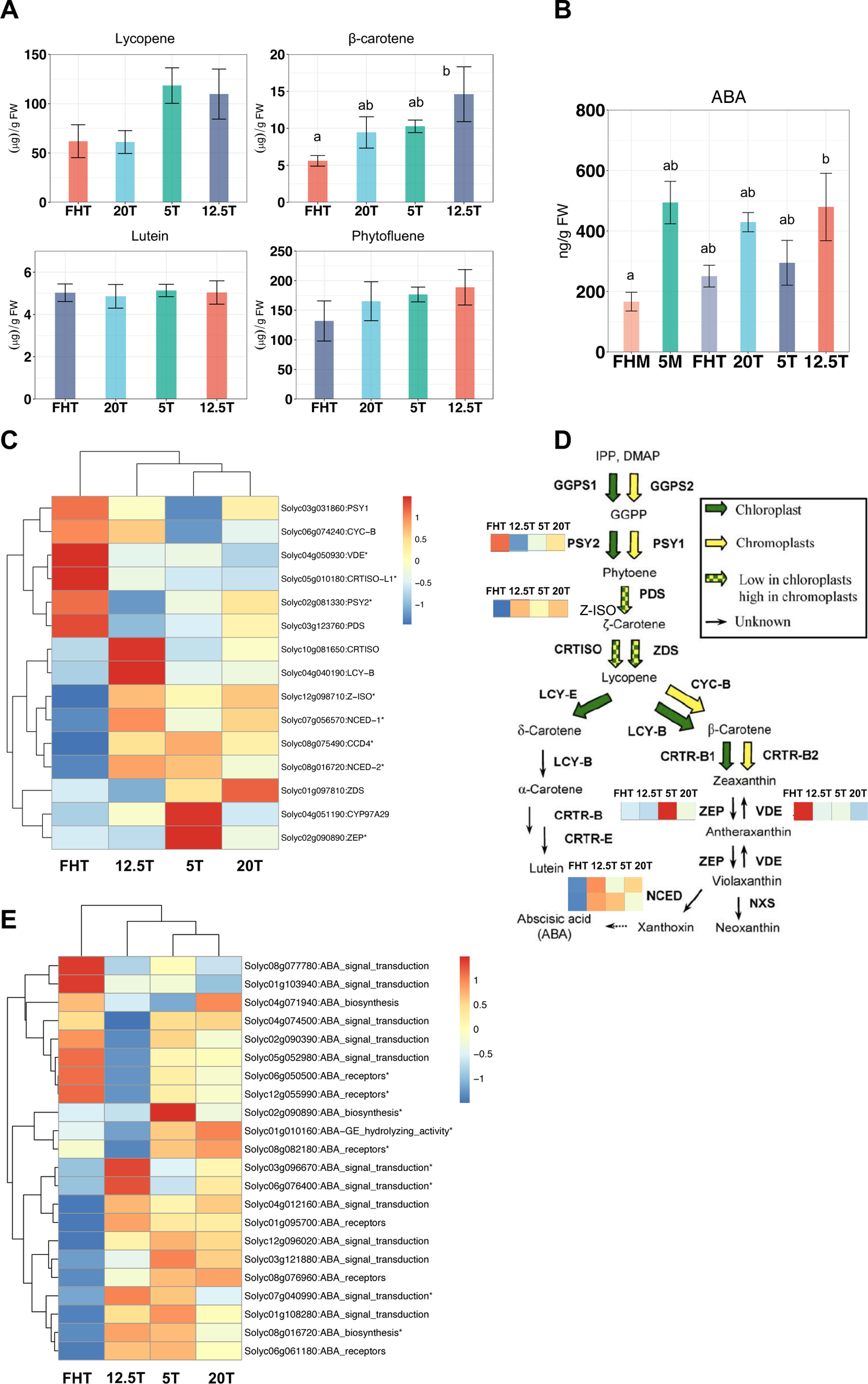
Fruit carotenoids and Abscisic acid (ABA). (**A**) carotenoid profiling includes lycopene, lutein, β-carotene and phytofluene. The error bar represents standard deviation of the mean. (**B**) Fruit ABA contents. The ‘5M’ only has two biological replicates, and others have three replicates. (**C**) Transcriptomic analysis of carotenoids biosynthesis and metabolic pathway. The Tukey multigroup tests were applied and asterisks were added only for the significant genes (*p* < 0.05), which applied to all gene expression heatmaps below. (**D**) Carotenoids biosynthesis pathway adapted from Galpaz *et al*., (2006) ^54^. Transcription heatmaps of the significantly DEGs were annotated on the side of the genes. (**E**) Transcriptomic analysis of ABA related genes.

Transcriptome analysis of the carotenoid-related pathway showed that *CRTISO*, *LCY-B*, *Z-ISO*, which are upstream β-carotene synthesis, were upregulated in ‘12.5T’ fruit (Figs. 4C-D). This may explain the high contents of β-carotene in ‘12.5T’. The enzymes encoded by *ZEP* and *VDE* inversely regulate β-carotene metabolism ^51^. *ZEP* was upregulated in ‘5T’ (2.28-fold vs. ‘FHT’), and *VDE* was upregulated in ‘FHT’ (15.03-fold vs. ‘12.5T’). However, post-transcriptional regulation of carotenogenic enzymes may potentially lead to non-linear connections between gene expression and carotenoid content.

ABA is produced downstream of the carotenoid biosynthesis pathway as a stress-responsive and ripening-related hormone (Fig. 4D). Fruit ABA increases from immature green to Turning, then decreases until red ripe ^45^. In accordance, all Turning fruit had higher ABA content than ‘FHM’ (Fig. 4B). With ‘FHM’ as the control, ‘12.5T’ fruit had more ABA (2.88-fold) accumulated than other Turning fruit (i.e., 1.51-fold in ‘FHT’, 2.58-fold in ‘20T’, 1.77-fold in ‘5T’; Fig. 4B). We examined the RNASeq data for connections between ABA and carotenoid gene expression. In ‘12.5T’ fruit, the two *NCED* isoforms that directly control ABA biosynthesis were expressed higher (*p* < 0.05) compared to ‘FHT’, i.e., (*NCED-*1; 3.94-fold and *NCED-*2;10.2-fold) (Fig. 4C). This may be an ABA-stress response activated at low temperature over a prolonged period, which tracked with higher levels of ABA in ‘12.5T’ fruit.

Other genes in the ABA biosynthesis and signaling pathway were examined (Fig. 4E). In ‘12.5T’, fruit ABA receptor genes were generally downregulated, and noticeably one ABA receptor, *SlPYL1* (*Solyc08g082180*) which has a known role in postharvest fruit ripening under low temperature ^52^, was significantly suppressed in ‘12.5T’. The *beta-glucosidase* (*Solyc01g010160*) can release free ABA by hydrolyzing ABA-GE ^53^, which was downregulated in ‘12.5T’. Interestingly, this is contrary to the high ABA content in that sample. In addition, expression of genes involved ABA signal transduction was remarkably high in ‘12.5T’ (Fig. 4E). These data indicate that ABA catabolism is complicated in 12.5°C fruit.

### 3.6 Postharvest fruit ethylene production and respiration rates

Ethylene and carbon dioxide production are characteristic of climacteric fruit ripening, and changes in the rate of production also serve as stress biomarkers for postharvest tomato ripening^13^. Ethylene production and respiration rates from MG until fruit ripening were depicted in Figs. 5A-B. The ethylene produced by ‘5M’ after rewarming was projected (dashed lines) onto the same timescale of the 20°C stored fruit, allowing comparisons between normal fruit ripening and stress-response-related ripening. First, total ethylene production under 20°C and 5°C rewarmed were similar (Table S8 and Fig. S11), indicating that chilling didn’t change the overall ethylene production but induced differences in production rates. Second, the rewarmed fruit had the characteristic intense burst of ethylene compared to normal ripening (20°C) (Figs. 5A and S11), indicating stress induced rapid ethylene accumulation. This sharp ethylene burst could trigger physiological decay of fruit quality compared to the normal ripening.

**Figure 5.**
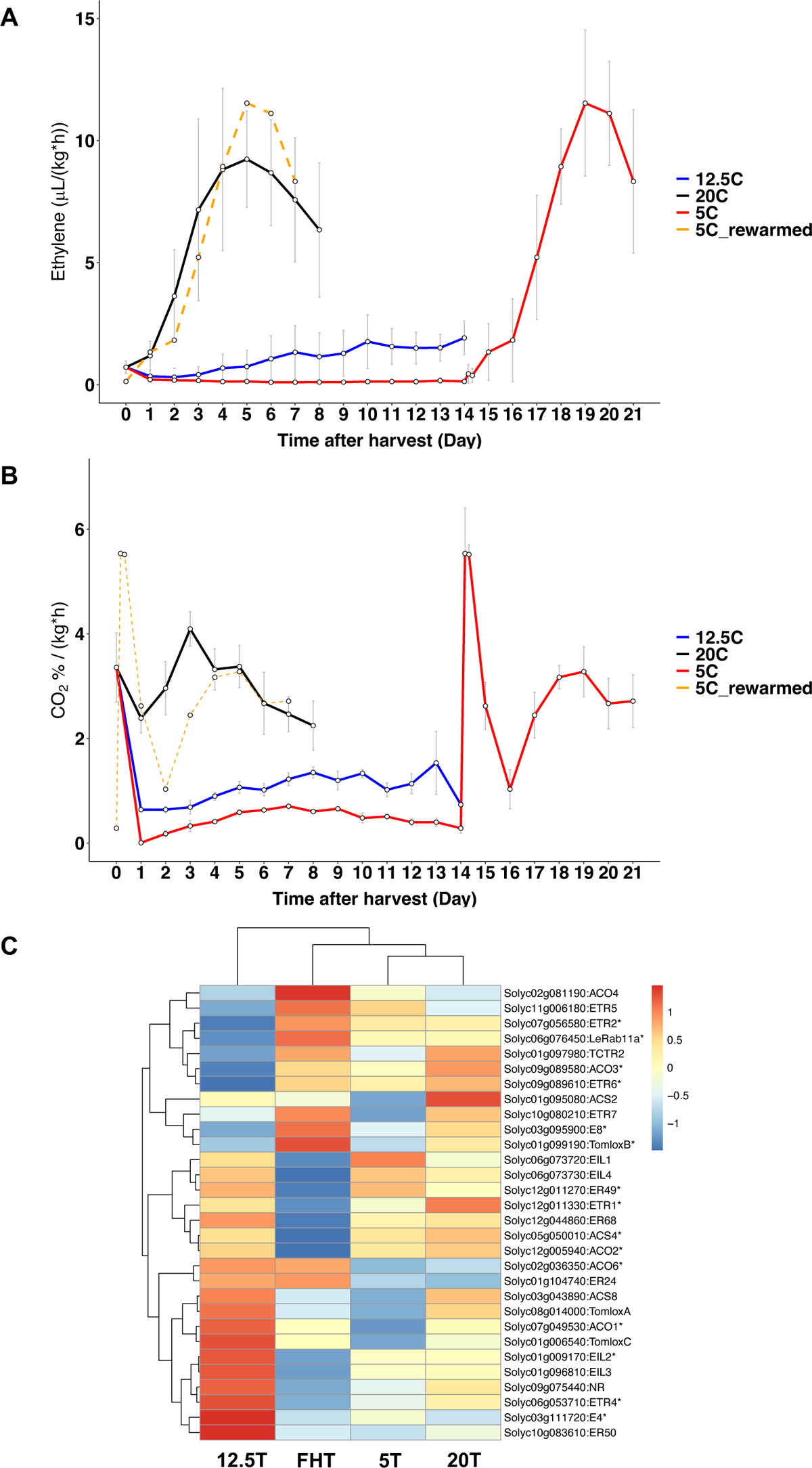
Ethylene and carbon dioxide production in relation to gene expression of the postharvest fruit. Ethylene production **(A)** and the CO2 levels **(B)** of the fruit harvested at the MG and stored at 20°C (black line), 12.5°C (blue line) and 5°C (red line) for 2 weeks and rewarmed to 20°C (red line). The rewarmed trend was also moved to the same x-axis scale (shown as the dashed orange line) to compare with ‘20C’. The error bar represents standard deviation of the mean. There are at least 6 biological replicates used in this assay. Tukey multigroup statistical tests were performed and shown in the Table S8. **(C)** Ethylene biosynthesis and related gene expression heatmap from RNASeq.

There were two peaks of respiratory activity in the rewarmed fruit (Fig. 5B). The first peak was likely the immediate stress response to increase metabolic activity for chilling injury recovery ^55^. The second peak occurred along with the ethylene burst at the day 18-19, which is the typical climacteric fruit respiratory burst ^56^. After day 4, the total CO_2_ production in the rewarmed fruit is close to that produced during normal ripening, indicated by the overlapping black and orange lines (Fig. 5B). In addition, the day 0 for all postharvest fruit showed the highest respiratory rates which could be due to the stress after harvest.

Strikingly, the 12.5°C fruit showed no obvious climacteric ripening peak of ethylene or CO_2_ over the 14-day storage, even though the fruit at this temperature underwent normal color development and quality changes ^5^. Furthermore, the 12.5°C fruit had reduced ethylene and CO_2_ total production compared to ‘20C’ and ‘5C_rewarmed’ during storage periods, though the fruit were stored for 14 days (Table S8).

The noteworthy question is if ethylene is the driving hormone for the apparent fruit ripening under 12.5°C. We, therefore, looked at the expression of genes involved in the ethylene pathway (Fig. 5C). The ‘12.5T’ fruit was still the most distinct group, as many ethylene pathway genes were upregulated in ‘12.5T’ but significantly downregulated in ‘FHT’, including *ACS4*, *ACO2*, *E4*, *ACO1*, *EIL2*, *ETR1* and *ETR4*. *ETR4* is a negative regulator of ethylene biosynthesis, and its repression results in faster fruit ripening ^57,58^. Of note is E4 because it is related to the transition between system 1 and system 2 ethylene biosynthesis and is strictly responsible for ethylene accumulation ^59^. In tomato ripening, expression for many ethylene genes, such as *ACO1*, *EIL2*, *ETR1*, *ETR4* etc., peaks at breaker, and then decreases at 10 days post breaker (https://bar.utoronto.ca/). The upregulation of these genes only happened in ‘12.5T’, suggesting that the 12.5°C storage may have a specific effect on the expression of these genes, possibly leading to a delay in their typical expression changes during fruit ripening. In summary, the ‘12.5T’ fruit had relatively low ethylene levels, no obvious ethylene system 2 peak, but high expression levels of some important ethylene-related genes. The mechanisms under those surprising findings may be related to the enhanced ABA in ‘12.5T’.

### 3.7 Fruit photosynthetic-related activity

The role of photosynthesis on tomato fruit ripening has been underestimated but was highlighted by the methylome and transcriptome data in this work. To determine if there was an association between the -omics data and the fruit photosynthetic markers, the delta absorbance (DA) index (I_DA_) was assessed by a non-destructive DA meter. The I_DA_ is the difference in absorbance between 670 nm and 720 nm, and chlorophyll *a,* the main chlorophyll in ripening tomato fruit, peaks at 660 nm ^60^. I_DA_ is highly correlated with fruit skin color and chlorophyll contents in tomato ^61^, and lower I_DA_ are recorded as the fruit ripens. In Figs. 6A and S12, the MG fruit have higher I_DA_ than Turning fruit. Specially, the ‘FHT’ is the highest compared to all postharvest Turning fruit, even though they have the same color characteristic. The gene expression is closely related to I_DA_, in which the ‘FHT’ is remarkably high in many photosynthetic genes (Fig. 6B). The dramatic changes in photosynthetic genes led to the next question, i.e., if the postharvest dark storage relates to the findings. To test this, we stored the MG fruit at 5°C under light or dark condition. The I_DA_ was assessed after 2 weeks. Surprisingly, the ‘5M’ under dark and light have no difference in their I_DA_. However, when compared to fresh harvest MG fruit, fruit I_DA_ in ‘5M’ under light is the same as ‘FHM’, while fruit I_DA_ in ‘5M’ under dark was significantly reduced (Fig. S12). The same result was found after repeating this experiment.

**Figure 6.**
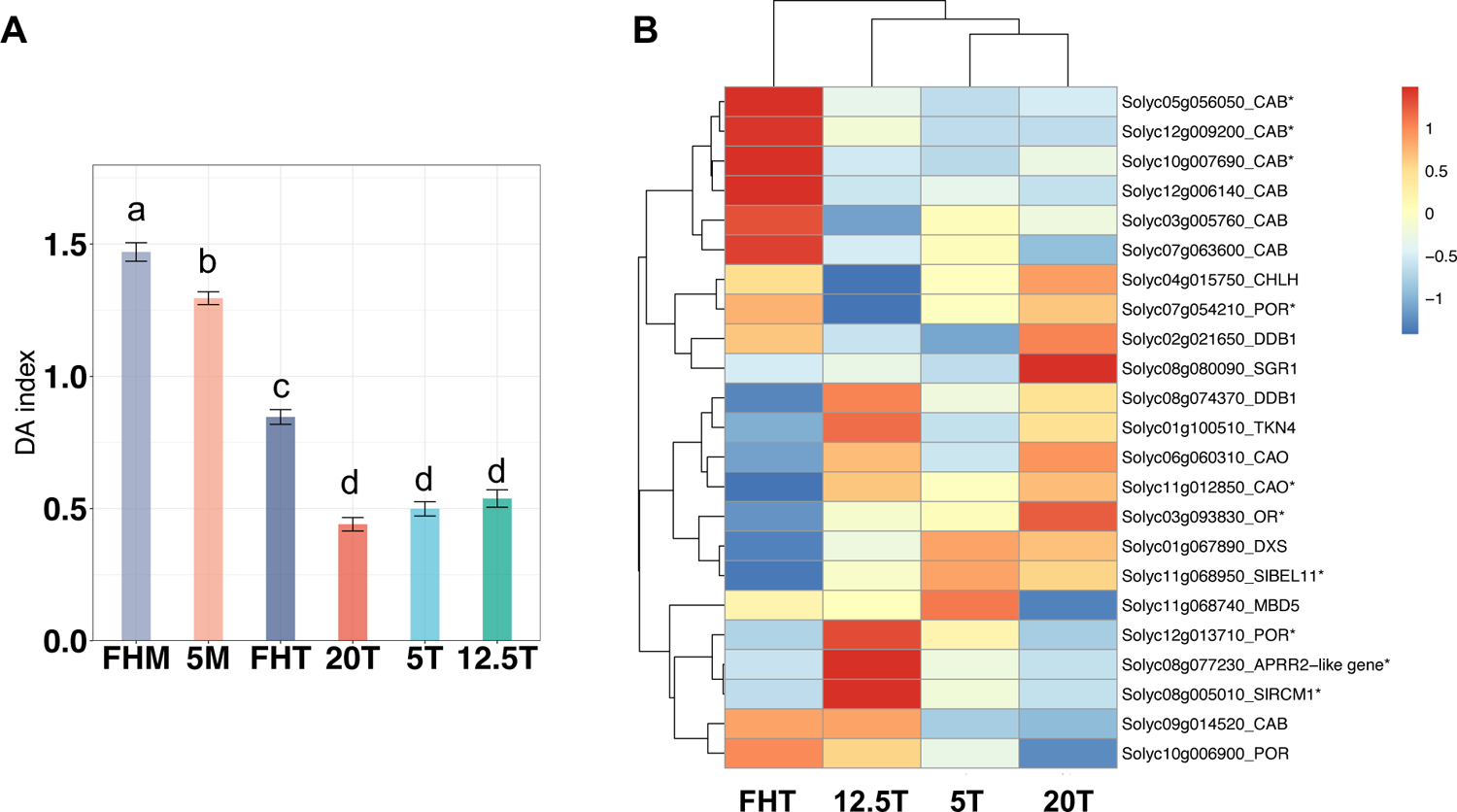
Postharvest tomato fruit IDA in relation to photosynthetic genes expression. (A) Postharvest fruit IDA. Each treatment includes 20 individual fruit as replicates. The Tukey multigroup tests were applied and the letters above each bar indicate the significance levels (*p* < 0.05). (B) Fruit photosynthetic related gene expression heatmap from RNASeq.

Correlative analyses between (1) the I_DA_ and gene expression, and (2) DNA methylation and transcription for photosynthetic genes were performed. The expression of four genes correlated (*p* < 0.05) with both the I_DA_, and changes in DNA methylation i.e., two *CAB* genes, *CAO* and *BEL11* (Figs. 6B and S13). Chlorophyllide *a* oxygenase (*CAO*) catalyzes chlorophyll *a* to chlorophyll *b*, and this gene was significantly downregulated in ‘FHT’ (Fig. S13). This may correlate to high I_DA_ (indicated chlorophyll *a* content) in ‘FHT’ (Fig. 6A). *CAB* are members of the <u>c</u>hlorophyll <u>a</u>/b <u>b</u>inding protein family, positively correlated with plant chlorophyll contents ^62^. *BEL11* is a negative regulator of fruit chloroplast development and induces photosynthesis related gene expression ^63^. ‘FHT’ fruit have high expression of *CAB* and reduced *BEL11*, which positively correlates to their high chlorophyll contents (Fig. 6).

### 3.8 Fruit ripening and quality related gene expression and methylation correlative analysis

We further examined the specific genes and transcriptional factors involved in ripening-to-senescence transition ^48^, auxin/IAA biosynthesis, cell wall metabolism and DNA methylation related. Analysis was on genes’ (1) transcriptional levels and (2) correlation between DNA methylation and gene expression.

The ripening gene expression heatmap (Fig. S7) clearly showed that the ‘12.5T’ fruit has a unique profile, with 32.7% genes (16 out of 49 genes) highly expressed in these fruit but not in fruit from the other treatments. While in ‘FHT’, 49.0% of genes are distinctively upregulated compared to other postharvest groups. The ‘12.5T’ distinct profile was also found in DNA methylation (Fig. S9) and auxin/IAA (Fig. S10) heatmaps. However, this trend was not consistent with cell wall related genes (Fig. S8), in which ‘12.5T’ and ‘5T’ have similar cell wall gene expression.

The DEGs of the fruit ripening and quality related pathways were examined for their correlation between gene expression and their gene associated DNA methylation levels. The genes with significant correlation were summarized in Tables 1-2. For the ripening related genes (Table 1), the *BEL11* plays a role in fruit ripening, and it negatively regulates chlorophyll biosynthetic genes^63^. *BEL11* is upregulated in ‘5T’ and ‘20T’ (Fig. S7), which may explain their reduced chlorophyll contents. *HDA1/3/5* are histone deacetylases, which control ripening by acting as transcriptional co-repressors ^64^; their differential expression in ‘12.5T’ may influence the ripening related genes expression. *ERF.C.3*, *HB1*, *MED25* and *WRKY17* are ethylene related, implying that postharvest handling affected ethylene regulation through DNA methylation. The two regions of *NAC-NOR*, master ripening regulator in tomato, have inverse expression-methylation correlation (*r* as 0.7466 and −0.6193), and its expression is remarkably high in ‘12.5T’.

**Table 1.**
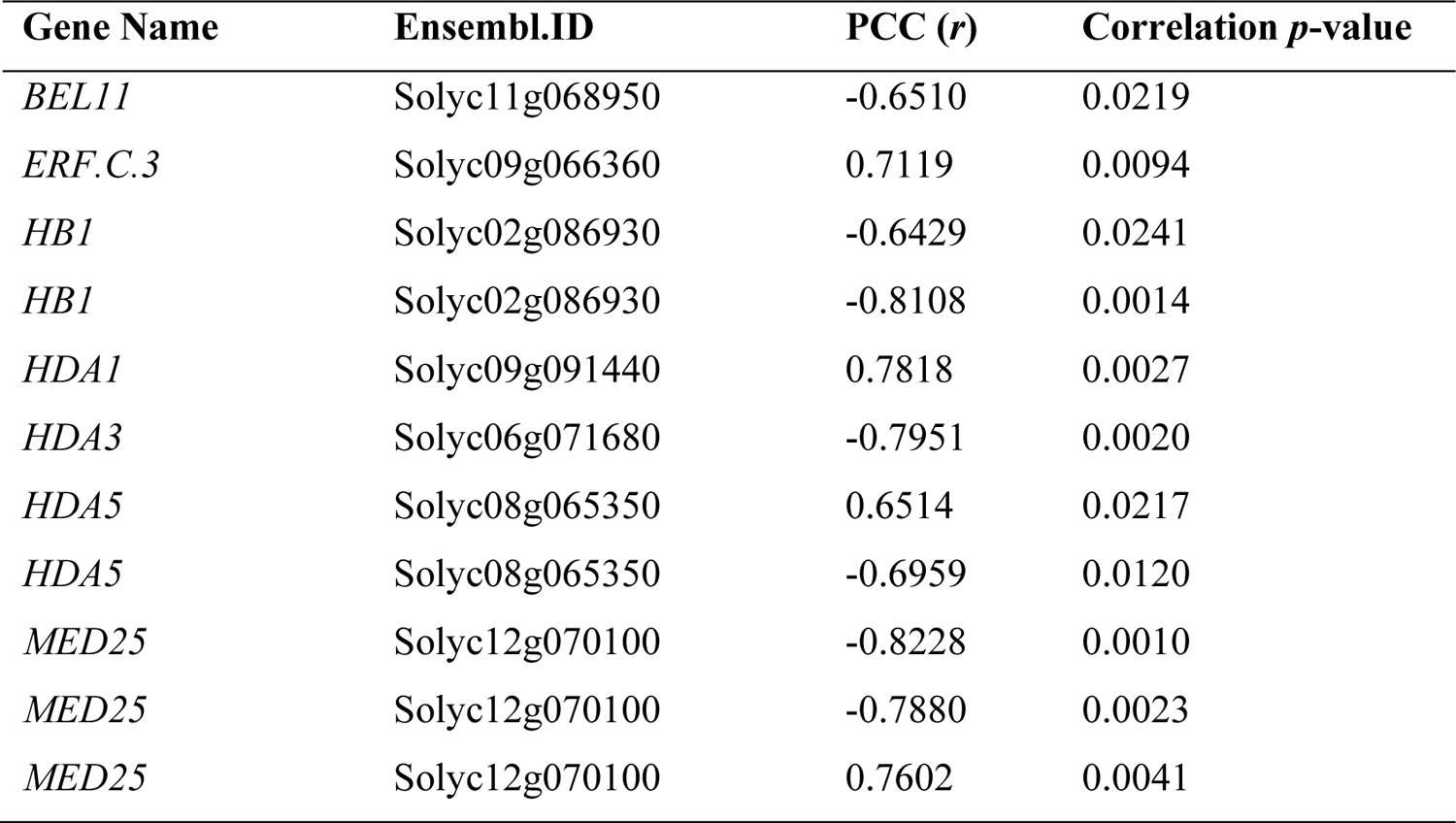
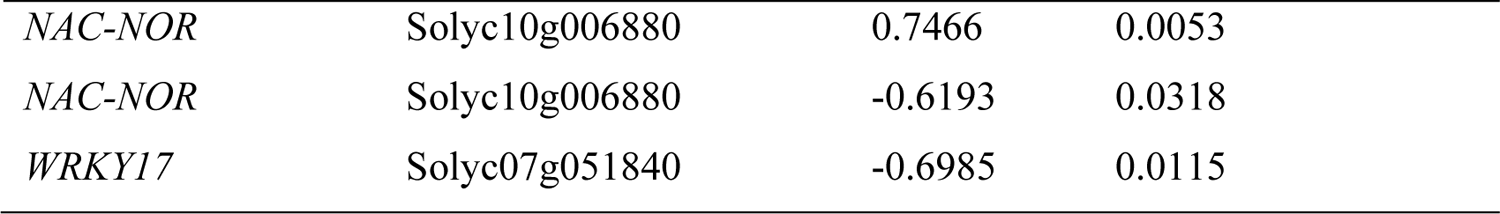
Ripening related DEGs (Fig. S7) with significant correlation between DNA methylation and gene expression. (The *p*-value 0.05 is as the threshold for both expression and correlation. The same genes may have multiple PCC due to multiple DNA methylation probes found in that gene associated regions.)

**Table 2.**
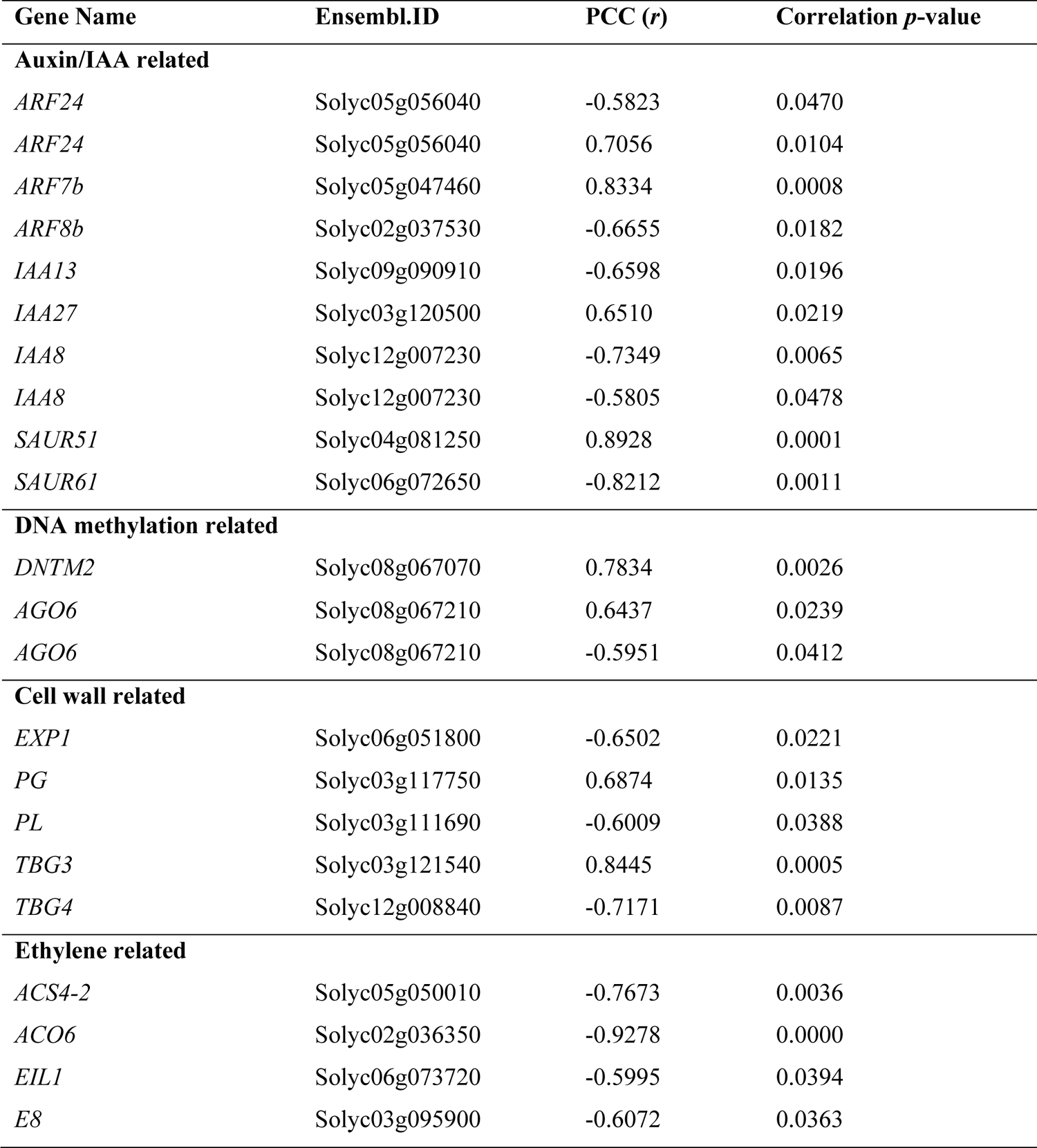
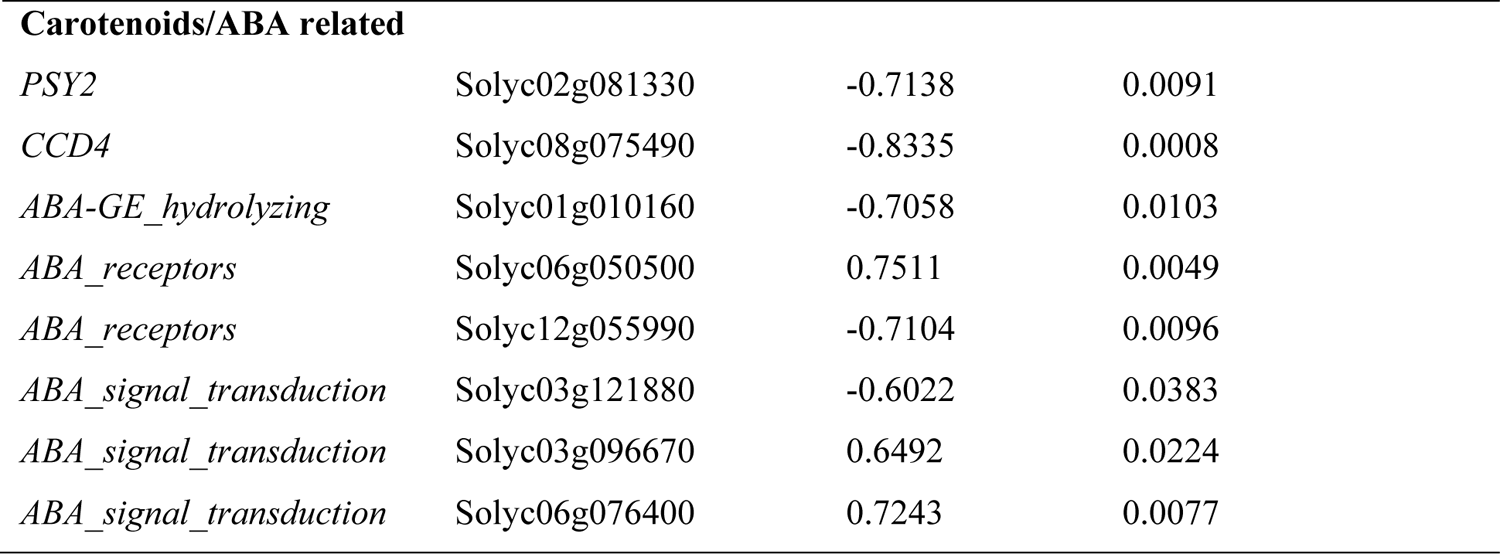
DEGs with significant correlation between DNA methylation and gene expression (According to Figs. S8-S10, 4B, 4E and 5C).

| Gene Name | EnsemblID | PCC (*r*) | Correlation *p*-value |
| --- | --- | --- | --- |
| **Auxin/IAA related** |  |  |  |
| *ARF24* | Solyc05g056040 | -0.5823 | 0.0470 |
| *ARF24* | Solyc05g056040 | 0.7056 | 0.0104 |
| *ARF7b* | Solyc05g047460 | 0.8334 | 0.0008 |
| *ARF8b* | Solyc02g037530 | -0.6655 | 0.0182 |
| *IAA13* | Solyc09g090910 | -0.6598 | 0.0196 |
| *IAA27* | Solyc03g120500 | 0.6510 | 0.0219 |
| *IAA8* | Solyc12g007230 | -0.7349 | 0.0065 |
| *IAA8* | Solyc12g007230 | -0.5805 | 0.0478 |
| *SAUR51* | Solyc04g081250 | 0.8928 | 0.0001 |
| *SAUR61* | Solyc06g072650 | -0.8212 | 0.0011 |
| **DNA methylation related** |  |  |  |
| *DNTM2* | Solyc08g067070 | 0.7834 | 0.0026 |
| *AGO6* | Solyc08g067210 | 0.6437 | 0.0239 |
| *AGO6* | Solyc08g067210 | -0.5951 | 0.0412 |
| **Cell wall related** |  |  |  |
| *EXP1* | Solyc06g051800 | -0.6502 | 0.0221 |
| *PG* | Solyc03g117750 | 0.6874 | 0.0135 |
| *PL* | Solyc03g111690 | -0.6009 | 0.0388 |
| *TBG3* | Solyc03g121540 | 0.8445 | 0.0005 |
| *TBG4* | Solyc12g008840 | -0.7171 | 0.0087 |
| **Ethylene related** |  |  |  |
| *ACS4-2* | Solyc05g050010 | -0.7673 | 0.0036 |
| *ACO6* | Solyc02g036350 | -0.9278 | 0.0000 |
| *EIL1* | Solyc06g073720 | -0.5995 | 0.0394 |
| *E8* | Solyc03g095900 | -0.6072 | 0.0363 |

| **Carotenoids/ABA related** |  |  |  |
| --- | --- | --- | --- |
| *PSY2* | Solyc02g081330 | -0.7138 | 0.0091 |
| *CCD4* | Solyc08g075490 | -0.8335 | 0.0008 |
| *ABA-GE_hydrolyzing* | Solyc01g010160 | -0.7058 | 0.0103 |
| *ABA_receptors* | Solyc06g050500 | 0.7511 | 0.0049 |
| *ABA_receptors* | Solyc12g055990 | -0.7104 | 0.0096 |
| *ABA_signal_transduction* | Solyc03g121880 | -0.6022 | 0.0383 |
| *ABA_signal_transduction* | Solyc03g096670 | 0.6492 | 0.0224 |
| *ABA_signal_transduction* | Solyc06g076400 | 0.7243 | 0.0077 |

## Discussion

Our objective was to investigate the impact of early harvest combined with postharvest storage at different temperatures on fruit DNA methylation. We also aimed to assess whether these postharvest conditions led to significant changes in gene expression in fruit ripening pathway and fruit physiology. Our transcriptomic and methylomic data revealed striking differences between fruit ripened after harvest and those ripened on the vine, irrespective of temperature storage. Notably, photosynthesis genes, were the primary determinants of this distinction. This is the first report that indicates substantial changes in the photosynthetic pathway in postharvest fruit. We also discovered that ‘12.5T’ fruit had the most distinctive DNA methylation and gene expression profiles, and it also displayed unique physiological traits, including carotenoids, ABA, and ethylene production.

Our work highlights significant changes in genes associated with ‘photosynthesis’ in postharvest fruit. The postharvest-stored fruit had reduced chlorophyll, supporting the clear distinction in the methylation status and expression of photosynthesis-associated genes. Fruit photosynthesis is a distinct process separate from leaf photosynthesis, primarily depending on CO_2_ refixation from respiration, as well as active but limited chloroplast activity ^65^. Many studies suggest that carbohydrates produced by fruit photosynthetic activity contribute to the energy and carbon required for synthesizing metabolites responsible for desirable fruit flavor attributes, maintaining O_2_ levels in the inner fruit tissue, and fueling seed development ^66–68^. These discussions on the importance of fruit photosynthesis have focused on green fruit with active chloroplasts. During ripening, chloroplast degradation and the development of chromoplasts, accompanied by a decline in chlorophyll and an increase in carotenoids, limit fruit photosynthesis ^69^. Our work is of note due to the upregulated photosynthetic transcriptional activity observed in Turning fruit on the vine compared to harvested fruit. This may underscore the significance of fruit photosynthetic activity during ripening. It’s worth noting that *SGR1,* a crucial gene in the process of tomato chlorophyll degradation ^70^, is uniformly expressed in all Turning fruit. *SGR1* is reported to be activated by fruit development and cold temperature ^71^, suggesting our postharvest treatments may not have direct effect on chlorophyll degradation. A recent study reported that fruit photosynthetic gene expression is upregulated in both green and ripened fruit under water stress when source capacity is constrained ^72^. This indicates a dynamic tradeoff between source and sink photosynthesis to support organ development.

Our work points to the strong effect of light on the methylome, transcriptome and chlorophyll levels of stored fruit compared to temperature and other stresses. Light is essential for fruit photosynthesis and chlorophyll synthesis ^73,74^. While chlorophyll captures light energy during photosynthesis, it may not always accurately predict photosynthetic activity. A proportional relationship between chlorophyll and photosynthetic rates may only occur under specific conditions and in certain plant tissue ^75^, although there is consistency in fruit chlorophyll contents, photochemical potential, and expression of photosynthesis related genes in Micro-Tom ^76^. Therefore, whether light has a direct effect on postharvest fruit photosynthesis requires more evidence. It has been suggested that CO_2_ evolution rates are higher in dark-stored tomato fruit than in those stored in the light, possibly due to reduced photosynthesis ^77^. Our data are suggestive and can be reinforced with measurements of net photosynthesis rates (change of CO_2_ levels), electron transport and Rubisco activities, in addition to chlorophyll contents, to accurately indicate postharvest fruit photosynthetic activity.

Beyond the possibility of photosynthesis occurrence, evidence for light influencing fruit metabolism is numerous. Light (1) enhances respiration, and induces an earlier onset climacteric ethylene peak, resulting in a shorter fruit shelf-life ^78^; (2) improves tomato nutritional quality and flavor ^79^; (3) controls fruit carotenoid development during ripening as an activation signal ^80^; (4) mediates signaling transduction associated with the methylation status of ripening genes’ promoters ^81^. Taken together, these studies support that restricted light, a common practice in postharvest handling, may contribute to quality reduction in postharvest fruit.

The low but non-chilling storage of ‘12.5T’ fruit leads to distinctive profiles of DNA methylation and gene expression patterns, and carotenoid levels. Most interestingly, the ‘12.5T’ fruit had no ethylene climacteric burst but relatively high levels of ABA. Our hypotheses are that (1) this low temperature storage without rewarming suppressed the normal climacteric peak, and (2) the complex hormone interplay of ethylene, ABA, IAA, GA or others collectively lead to this biological ripening process ^82^. Remarkably, since ABA is proposed to act upstream of ethylene in tomato ripening ^19^, an uncoupled ripening process may occur between ABA and ethylene in ‘12.5T’. Ethylene production in ‘12.5T’ may lag behind ABA production, leading to the unique molecular regulation observed in this work. Moreover, while there are reports on how chilling temperatures inhibit ripening and alter hormone interactions, there is a lack of studies investigating the effects of low but non-chilling temperatures ^83–85^. In addition, ‘12.5T’ showed inconsistent results in gene expression validation using qRT-PCR, but there was high similarity in results between the two methods, i.e., RNASeq and qRT-PCR, in all other groups (Fig. S15). These conflicting results indicate that pre-harvest environments across growth seasons, significantly affect fruit gene expression after storage at 12.5°C ^86^. This effect may be magnified because of the extended developmental program of these fruit, and near the chilling temperature threshold, many chilling-related biological processes may be easily triggered.

We conducted a comparative study using two fruit stages, i.e., ‘Mature green’ and ‘Turning’. ‘Turning (T)’ is the ripening stage we selected for sampling and subsequent studies, because (1) both the fruit stored at 5°C followed by rewarming and the fruit at 12.5°C consistently reached the ‘Turning’ stage, but not red ripe, and (2) in ‘Micro-Tom’, Turning corresponds to the ‘Pink’ that is the stage just before red ripe in conventional tomato cultivars ^5,87^. Studying the ‘Turning’ stage enables us to capture differential gene regulation associated with ripening and quality before fruit senescence which begins at red ripe. We compared postharvest fruit to the fresh harvest fruit with identical color attributes, which we used as a proxy for fruit developmental stage; however, there is a disconnect between the physiological and chronological age of fruit ripened postharvest. The ‘12.5T’ fruit that took the longest time to ripen from MG to Turning had the highest methylation levels among all the Turning fruit (Fig. 1C). The fruit industry commonly uses color or other quality traits to define produce age. Instead, our data implied that methylome and transcriptome indicated age may be more accurate than cellular or chronological age ^88^. These fruit genomic molecular fingerprints could potentially serve as quality biomarkers for differentiating fruit internal quality parameters from external appearance, therefore contributing to a reduction in postharvest waste in the future.

For our -omic studies, we used bulk sequencing, which indicates the average percentage of methylation and the average levels of gene expression across millions of cells. Correlative analysis between methylation and expression was established for known ripening genes, and the genes with significant correlation were highlighted (Tables 1-2 and Fig. S13). This information is important for crop improvement through epigenome engineering ^89^. It is noteworthy that although we used low (3∼4 X) coverage of the tomato genome by bisulfite sequencing, the biological replicates remained consistent, and the methylation percentages closely aligned with results from a WGBS study using single-base resolution ^20^. Our study, along with the work of Crary-Dooley *et al*., ^90^ collectively supports the feasibility and reliability of low-coverage sequencing.

In conclusion, the analysis of -omics and physiological data in this work revealed that early harvest and storage have an impact on fruit ripening quality, hormone composition, and the transcriptome. Variations in many of these biological entities are closely associated with DNA methylation, as demonstrated by the expression-methylation correlations observed in many ripening genes. The integrative analysis of gene expression and DNA methylation correlation tests across multiple ripening and quality pathways pinpointed postharvest biomarker genes for future studies on tomato postharvest biology.

## Acknowledgments

JZ, BC and KA received Fellowships from the University of California Davis Graduate Group of Horticulture & Agronomy, and Henry A. Jastro Graduate Research Awards; KS and KL thank Postharvest Technology Innovation Center, Ministry of Higher Education, Science, Research and Innovation, Bangkok, Thailand; Work in DB’s lab was supported by the USDA National Institute of Food and Agriculture, Hatch Project CA-D-PLS-2404-H, and a UC Davis ADVANCE Scholar Fellowship. The authors thank Emma Shipman, Dave Tseng, Yichao Xu, Chiu-Ling Yang and Po-kai Huang for their critical reading of this paper, and the technical support of undergraduate interns Roshmund Romero and Annika Uemura.

## Contributions

JZ-conceptualization; methodology; formal analysis; led the bioinformatics analysis, investigation; validation; writing-original draft, review& editing; SZ-bioinformatics analysis; review& editing; BC-conceptualization; review& editing; KS-methodology (carotenoids assay); review& editing; KL-supervision and funding acquisition; review; KA-review & editing; DB-conceptualization; funding acquisition; methodology; project administration; resources; supervision; writing-original draft, review& editing.

## Data availability

The RNASeq and WGBS sequencing data has been uploaded to Sequence Read Archive (SRA) (http://www.ncbi.nlm.nih.gov/), and the BioProject ID is PRJNA1026769.

