## supplementary files for "Integrative analysis of the methylome and transcriptome of tomato fruit (*Solanum lycopersicum* L.) induced by postharvest handling"

### List of tables and figures in supplementary information

Supplementary Table S1. Bisulfite Sequencing statistic by Bismark and RNASeq reads statistics

Supplementary Table S2. Differential methylated genes (DMGs) expressed in fruit

Supplementary Table S3. DMGs Geno ontology functional analysis

Supplementary Table S4. Differential expressed genes (DEGs) summary

Supplementary Table S5. DEGs gene ontology functional analysis

Supplementary Table S6. Hub genes from WGCNA

Supplementary Table S7. Summary of DEGs with significant correlation between DNA methylation and gene expression

Supplementary Table S8. Ethylene and $CO_2$ assay total production rates and statistical test

Supplementary Figure S1. Postharvest treatment experimental design

Supplementary Figure S2. Methylation percentage in CG, CHG and CHH context

Supplementary Figure S3. Circular plot of methylome

Supplementary Figure S4. Context specific DMR annotations

Supplementary Figure S5. Weighted gene co-expression network analysis (WGCNA).

Supplementary Figure S6. GO figure output of Weighted gene co-expression network analysis (WGCNA)

Supplementary Figure S7. Transcriptomic analysis in fruit ripening related genes and transcriptional factors

Supplementary Figure S8. Transcriptomic analysis in fruit cell wall pathway

Supplementary Figure S9. Transcriptomic analysis in DNA methylation related genes

Supplementary Figure S10. Transcriptomic analysis in auxin/IAA related genes

Supplementary Figure S11. Ethylene production fitting curves

Supplementary Figure S12. Boxplot of the postharvest fruit DA index

Supplementary Figure S13. Photosynthetic genes with significant correlation and difference in expression

Supplementary Figure S14. Transcriptomic analysis by KEGG annotation

Supplementary Figure S15. qRT-PCR validation of the RNASeq DEGs

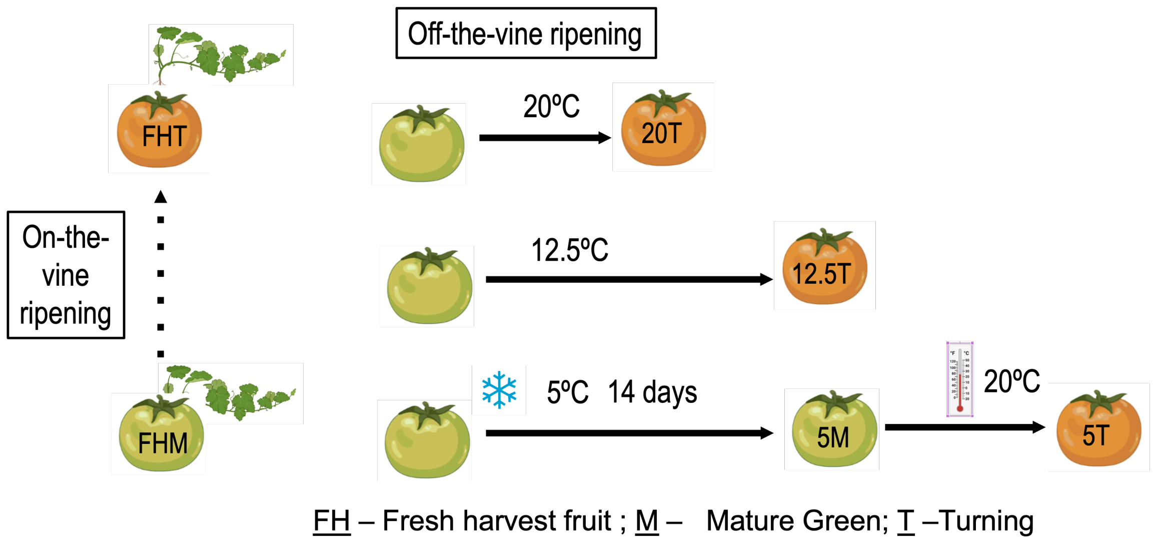


Figure S1 Postharvest fruit experimental design (adapted from Zhou *et al.*, 2021 [1]). The time harvested fruit taken from 'MG' to 'T' is indicated as the relative length of the black solid lines.

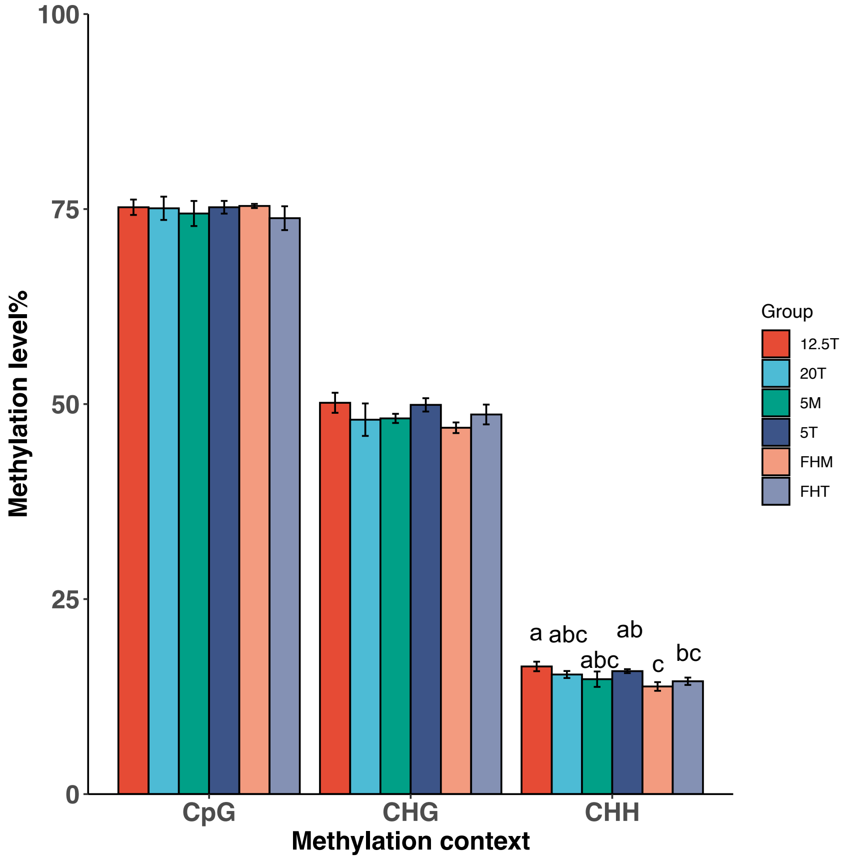


Figure S2 Genome methylation in tomato fruit at 'Mature green' (M) and 'Turning' (T) after fresh-harvesting (FH) or storage at different temperatures indicated. Methylation of the '12.5T' fruit was significantly higher than 'FHM' fruit ($p < 0.05$) in the CHH context.

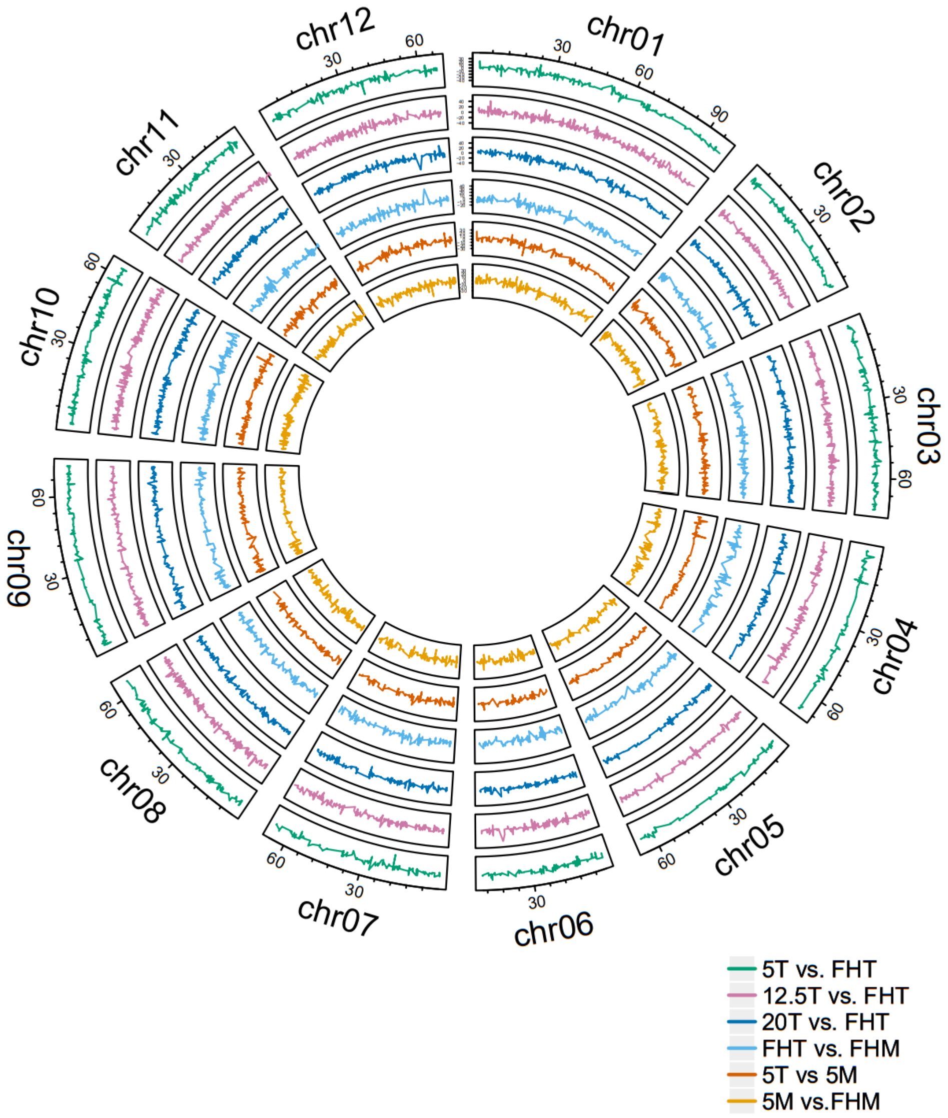


Figure S3 Circular plot of the DNA global methylation difference in each comparison. For each sector, the x-axis represents the methylated cytosine position (in Mb) in the chromosome, and the y-axis represents the methylation difference.

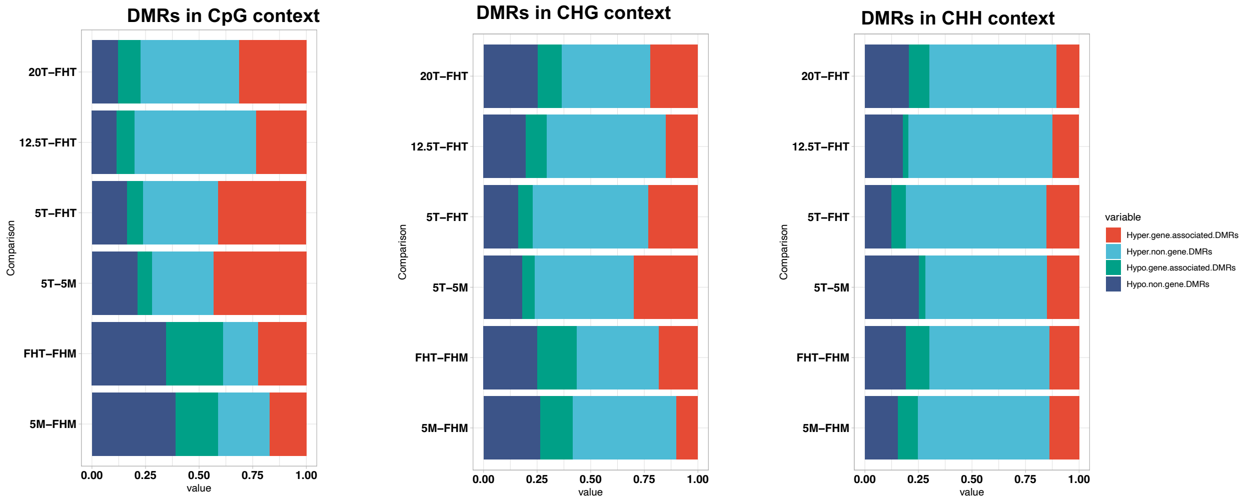


Figure S4 Proportion of differentially methylated regions (DMRs) in each comparison that are hypo and hyper-methylated, and if they are in gene and non-gene regions.

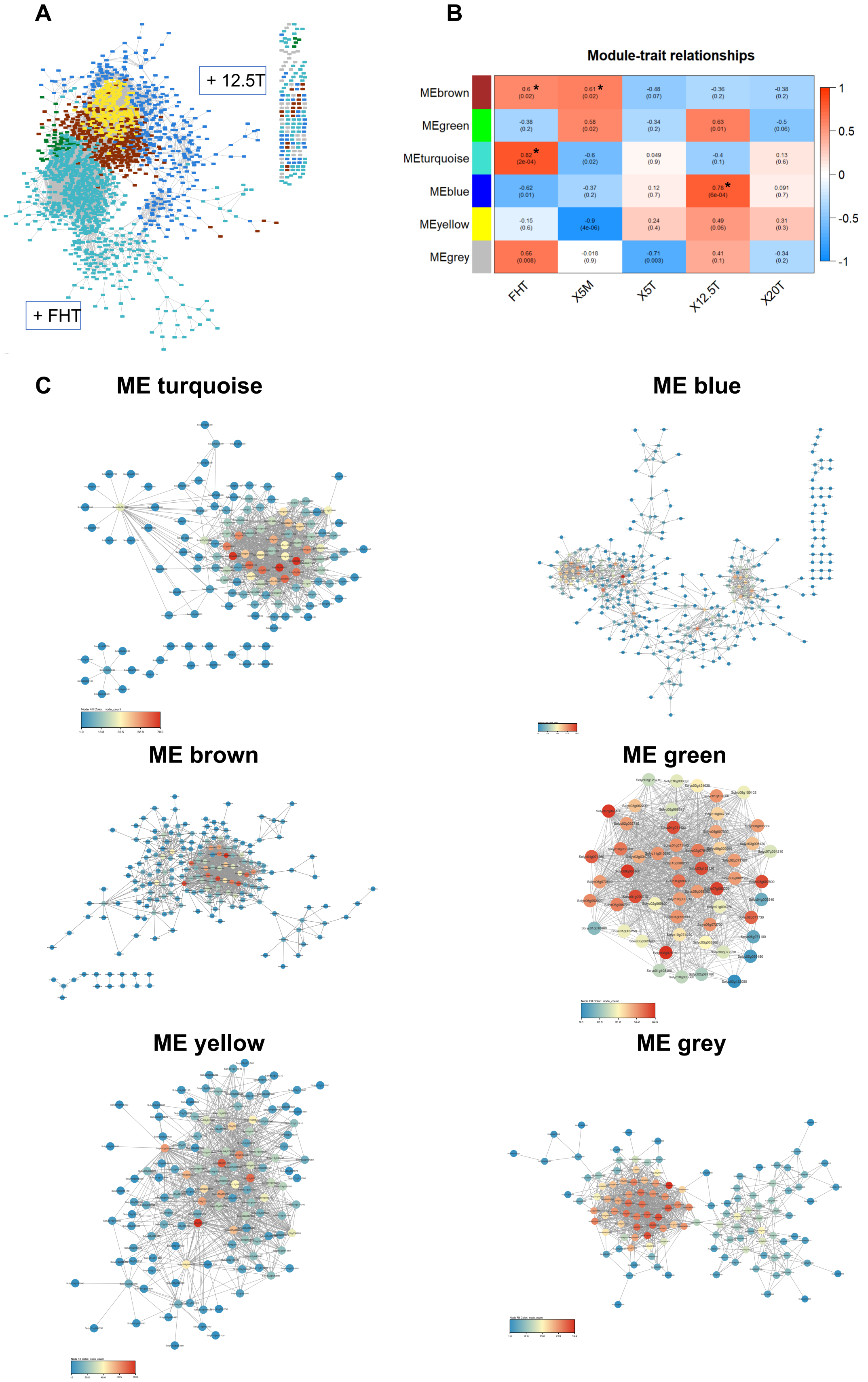
A
+ 12.5T
+ FHT
B
Module-trait relationships
MEbrown
MEgreen
MEturquoise
MEblue
MEyellow
MEgrey
0.6 * (0.02)
0.61 * (0.02)
-0.48 (0.07)
-0.36 (0.2)
-0.38 (0.2)
-0.38 (0.2)
0.58 (0.02)
-0.34 (0.2)
0.63 (0.01)
-0.5 (0.06)
0.82 * (2e-04)
-0.6 (0.02)
0.049 (0.9)
-0.4 (0.1)
0.13 (0.6)
-0.62 (0.01)
-0.37 (0.2)
0.12 (0.7)
0.78 * (6e-04)
0.091 (0.7)
-0.15 (0.6)
-0.9 (4e-06)
0.24 (0.4)
0.49 (0.06)
0.31 (0.3)
0.66 (0.008)
-0.018 (0.9)
-0.71 (0.003)
0.41 (0.1)
-0.34 (0.2)
FHT
X5M
X5T
X12.5T
X20T
1
0.5
0
-0.5
-1
C
ME turquoise
ME blue
ME brown
ME green
ME yellow
ME grey

Figure S5 Weighted gene co-expression network analysis (WGCNA). (A). Global view of all clusters explored in this study. The different colors represent different gene clusters. (B). A heatmap of eigengene expression across treatment groups and the module-trait correlations of five samples (‘FHT’, ‘5T’, ‘12.5T’, ‘20T’, and ‘5M’). The top value in each block represents the Pearson’s correlation coefficient (*r*) calculated between the module and the treatment group, and the lower value represents the corresponding *p-value* (* = $p < 0.001$ in at least one turning group). (C). Network of genes in top 1000 connectivity identified in each ME. The color of each node represents the node degree.

### ME turquoise

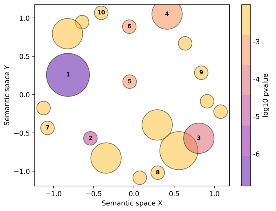


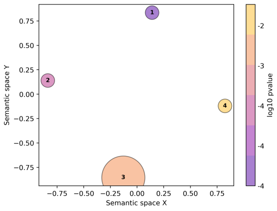


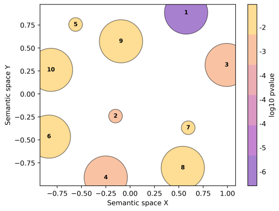


1. response to hydrogen peroxide
2. protein complex oligomerization
3. branched-chain amino acid biosynthetic process
4. response to heat
5. protein folding
6. positive regulation of superoxide dismutase activity
7. pollen maturation
8. Lewis a epitope biosynthetic process
9. polyketide biosynthetic process
10. positive regulation of defense response to insect

1. cytoplasm
2. chloroplast stroma
3. respiratory chain complex II
4. chloroplast

1. protein self-association
2. naringenin-chalcone synthase activity
3. transition metal ion binding
4. dihydroxy-acid dehydratase activity
5. beta-amylase activity
6. serine-tRNA ligase activity
7. oxidoreductase activity
8. 9-cis-epoxycarotenoid dioxygenase activity
9. phosphatidylinositol-4,5-bisphosphate 5-phosphatase activity
10. GTPase activity

### ME blue

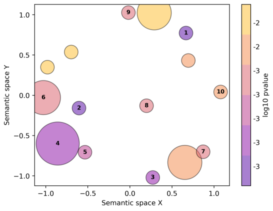


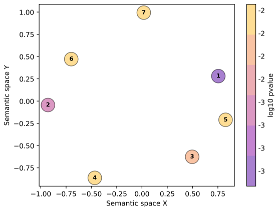


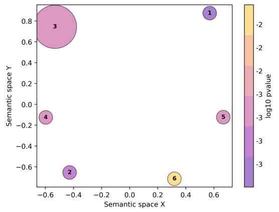


1. fatty acid biosynthetic process
2. regulation of nitrogen utilization
3. response to fungus
4. negative regulation of cell cycle
5. sequestering of actin monomers
6. regulation of auxin mediated signaling pathway
7. detection of ethylene stimulus
8. nucleus organization
9. DNA ligation involved in DNA repair
10. fruit ripening

1. chaperonin-containing T-complex
2. nucleolus
3. nuclear pore central transport channel
4. cell cortex
5. acetyl-CoA carboxylase complex
6. chloroplast thylakoid
7. chloroplast stroma

1. actin monomer binding
2. isocitrate dehydrogenase (NADP+) activity
3. DNA ligase activity
4. oxidoreductase activity, acting on the aldehyde or oxo group of donors, disulfide as acceptor
5. 1-aminocyclopropane-1-carboxylate oxidase activity
6. dehydropipecolic acid reductase

### ME brown

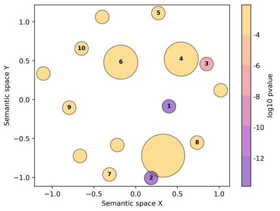


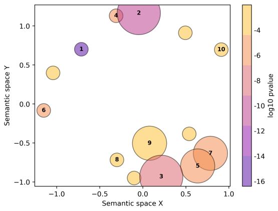


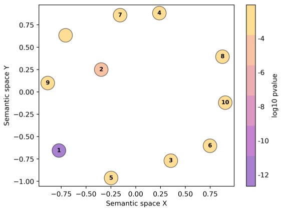


1. photosynthesis
2. translation
3. photosynthetic electron transport in photosystem I
4. photosynthesis, light harvesting in photosystem I
5. response to cytokinin
6. ribosomal small subunit assembly
7. starch biosynthetic process
8. maturation of SSU-rRNA
9. retrograde vesicle-mediated transport, Golgi to endoplasmic reticulum
10. brassinosteroid mediated signaling pathway

1. chloroplast thylakoid membrane
2. ribosome
3. photosystem II
4. thylakoid
5. cytosolic small ribosomal subunit
6. chloroplast envelope
7. cytosolic large ribosomal subunit
8. photosystem I reaction center
9. photosystem II oxygen evolving complex
10. chloroplast stroma

1. structural constituent of ribosome
2. NAD binding
3. catalytic activity
4. AU-rich element binding
5. transporter activity
6. aminoacyl-tRNA hydrolase activity
7. 5S rRNA binding
8. glyceraldehyde-3-phosphate dehydrogenase (NAD+) (phosphorylating) activity
9. chlorophyll binding
10. fructose-bisphosphate aldolase activity

### ME green

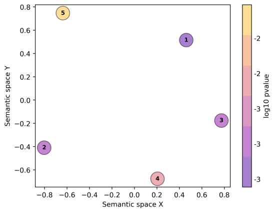


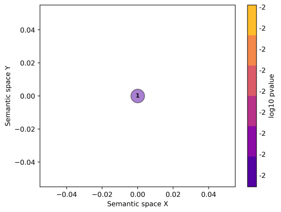


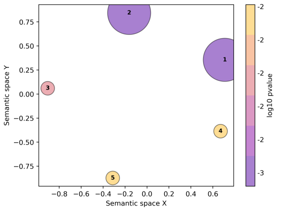


1. cyclic nucleotide metabolic process
2. protein heterotrimerization
3. ethanolamine metabolic process
4. threonine biosynthetic process
5. L-cystine transport

1. chloroplast stroma

1. aspartate-semialdehyde dehydrogenase activity
2. 2',3'-cyclic-nucleotide 3'-phosphodiesterase activity
3. L-cystine transmembrane transporter activity
4. protochlorophyllide reductase activity
5. NADP binding

### ME yellow

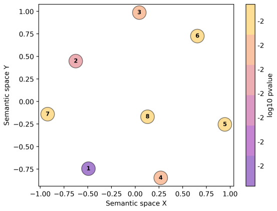


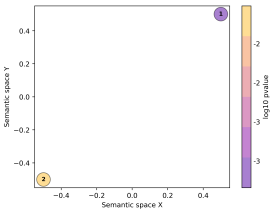


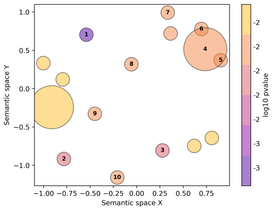


1. photosynthesis
2. starch biosynthetic process
3. regulation of tetrapyrrole metabolic process
4. fatty acid alpha-oxidation
5. UDP-galactose transmembrane transport
6. positive regulation of nuclear-transcribed mRNA poly(A) tail shortening
7. oxidative photosynthetic carbon pathway
8. lysine biosynthetic process via aminoadipic acid

1. photosystem I
2. cytoplasm

1. catalytic activity
2. calcium-dependent phospholipid binding
3. galactolipase activity
4. 3-oxo-arachidoyl-CoA synthase activity
5. 3-oxo-cerotoyl-CoA synthase activity
6. 3-oxo-lignoceronyl-CoA synthase activity
7. very-long-chain 3-ketoacyl-CoA synthase activity
8. UDP-glucosyltransferase activity
9. magnesium-protoporphyrin IX monomethyl ester (oxidative) cyclase activity
10. UDP-galactose transmembrane transporter activity

### ME grey

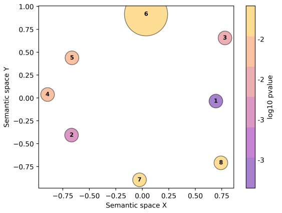


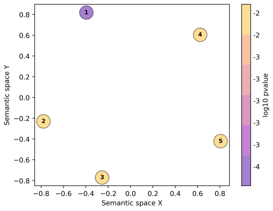


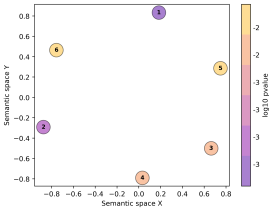


1. cellular oxidant detoxification
2. calcium-mediated signaling
3. carbohydrate transport
4. induced systemic resistance, ethylene mediated signaling pathway
5. positive regulation of transcription by RNA polymerase I
6. protein K69-linked ufmylation
7. microtubule-based process
8. response to cold

1. chloroplast stroma
2. mitochondrial proton-transporting ATP synthase, stator stalk
3. t-UTP complex
4. plant-type cell wall
5. extracellular exosome

1. intramolecular lyase activity
2. fatty acid binding
3. MAP kinase activity
4. aminomethyltransferase activity
5. catalytic activity
6. copper ion binding

Figure S6 GO figure output of Weighted gene co-expression network analysis (WGCNA) in each ME. The GO output is presented by three categories, i.e., biological progress, cellular component, and molecular function.

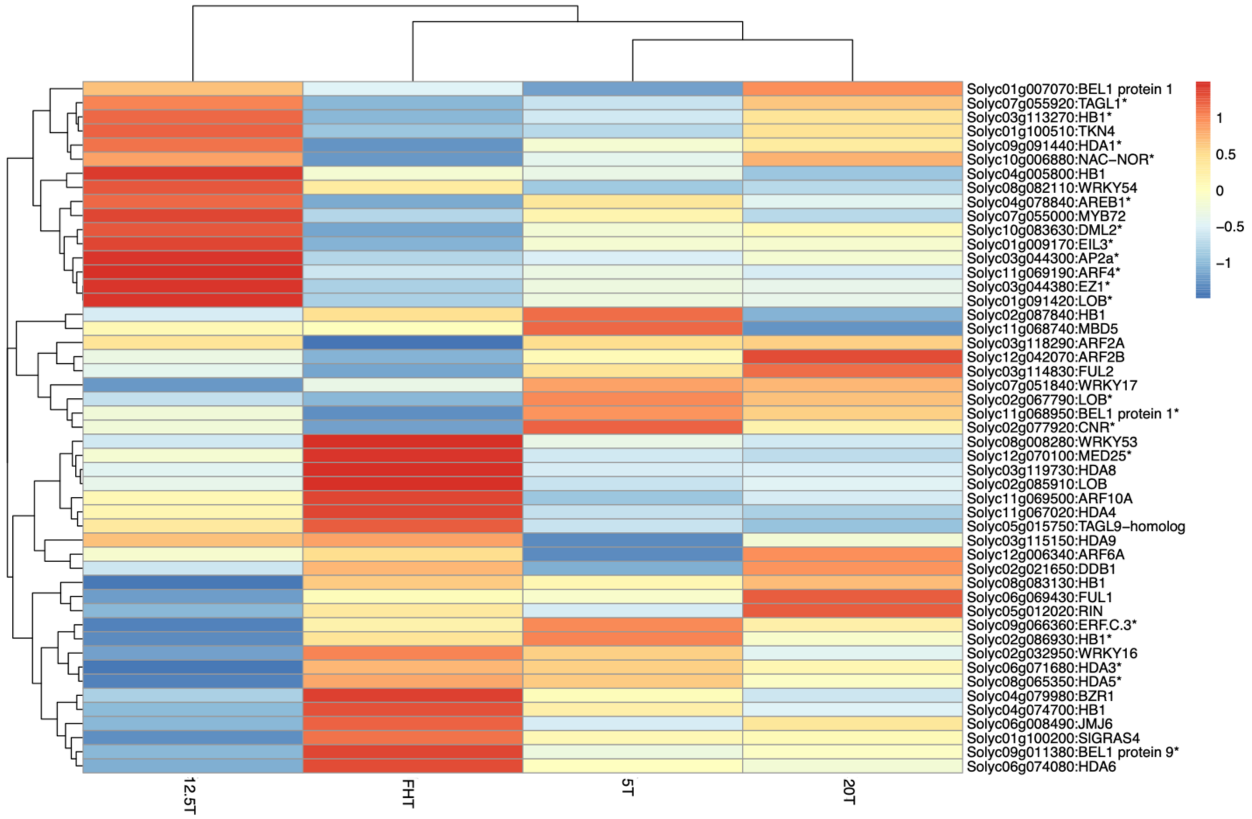


Figure S7 Transcriptomic analysis in fruit ripening related genes and transcriptional factors.

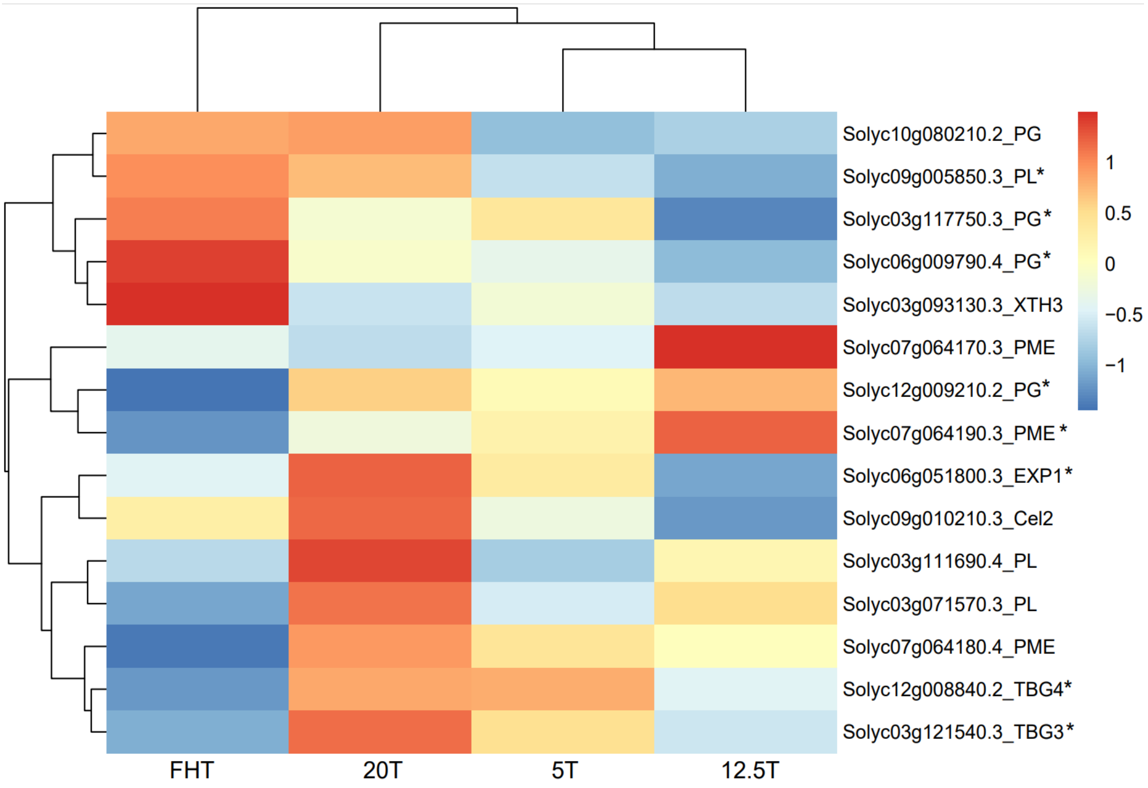


Figure S8 Transcriptomic analysis in fruit cell wall pathway.

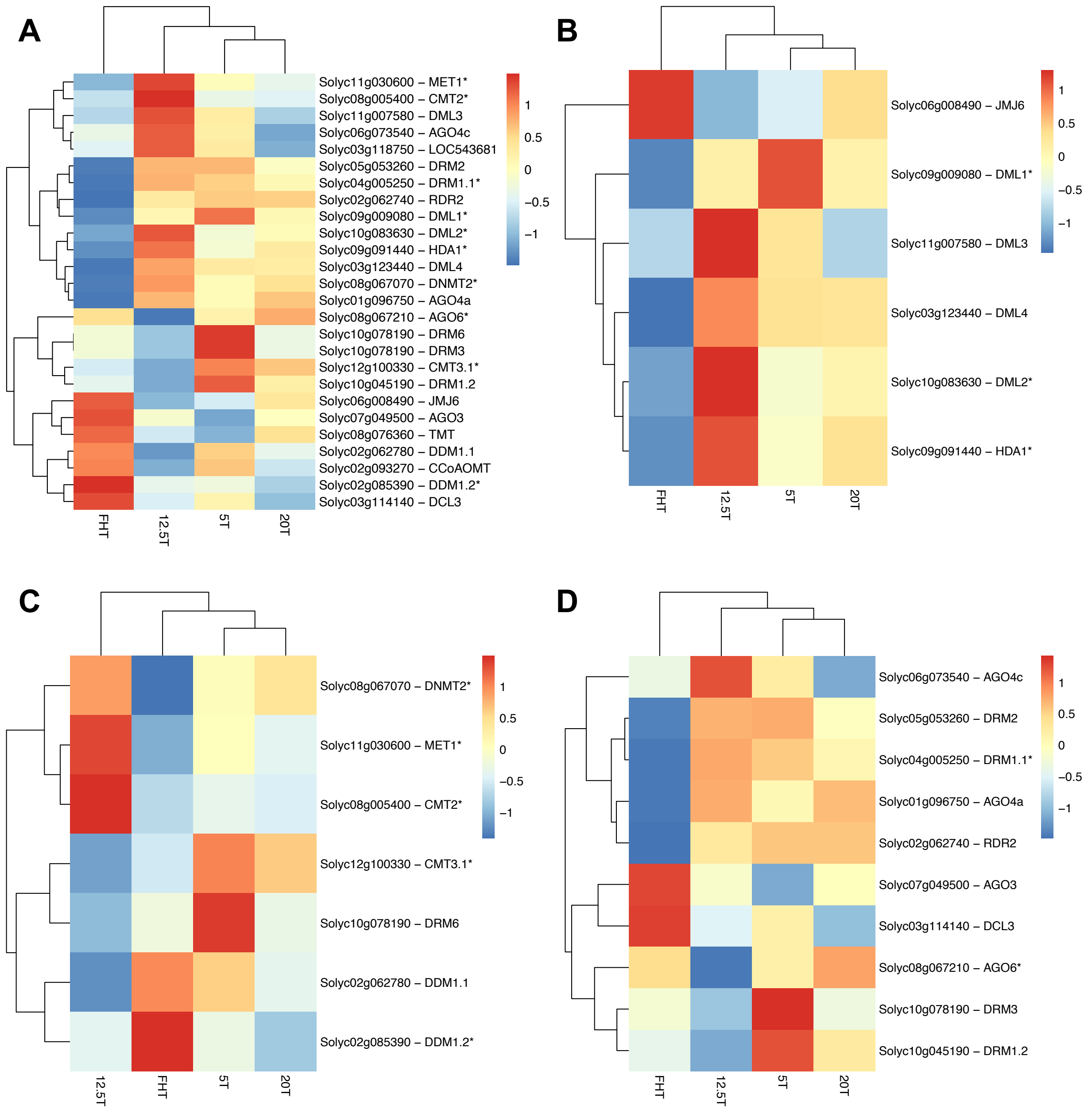


Figure S9 Transcriptomic analysis in DNA methylation related pathway. (A). Overall DNA methylation related genes expressed in the tomato fruit, which includes (B). Demethylation related, (C) Methylation maintenance, (D). RNA-directed DNA methylation (RdDM) pathway related genes.

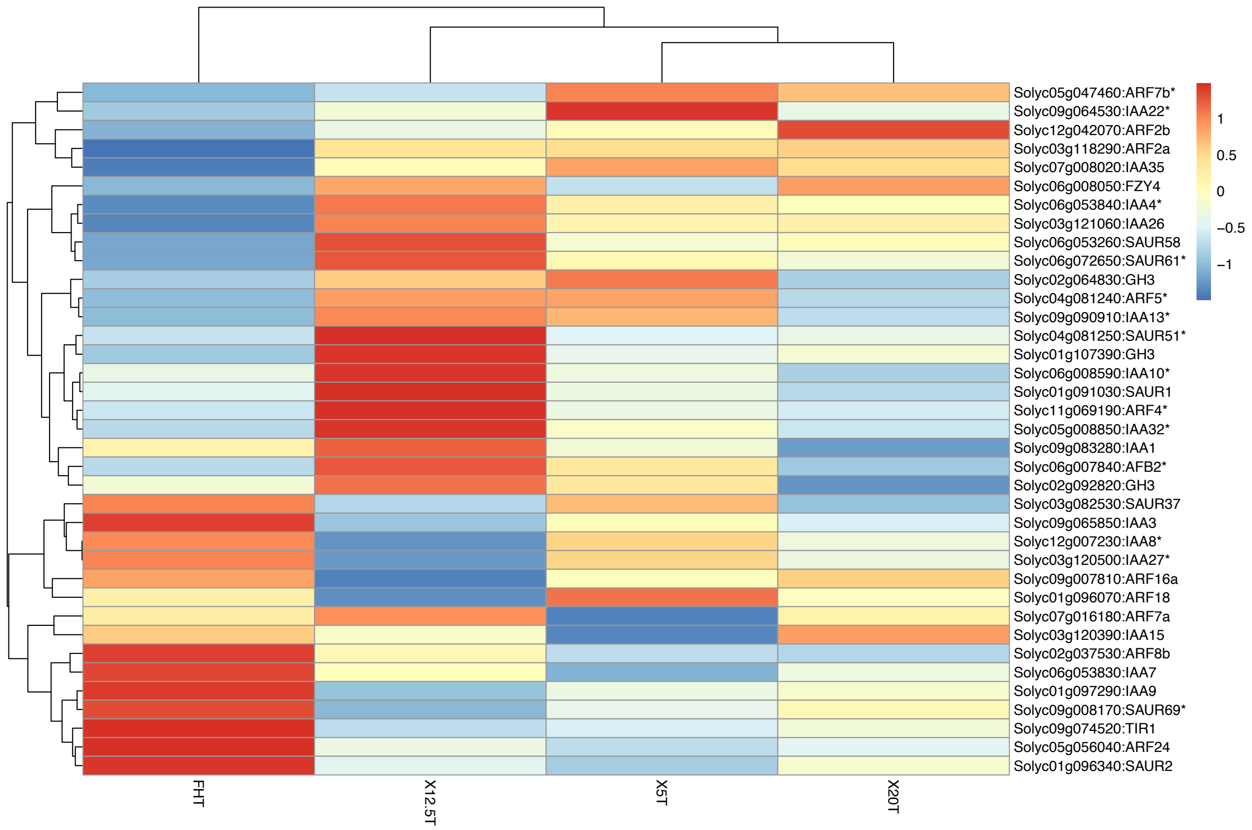


Figure S10 Transcriptomic analysis in auxin/IAA related genes.

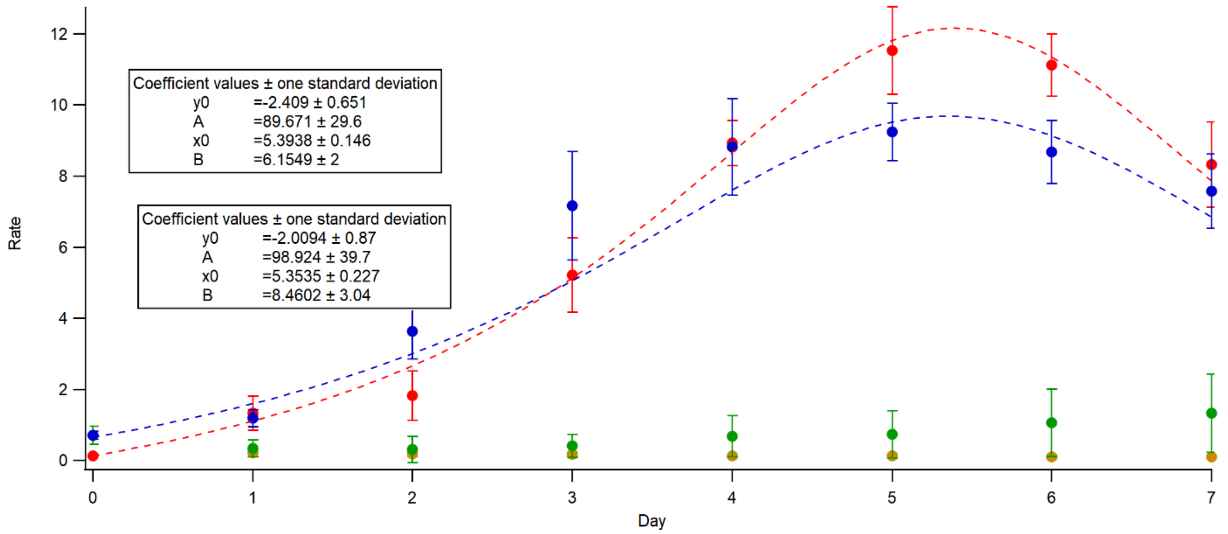


Figure S11 Ethylene production fitting curves. Ethylene of the fruit harvested at the MG and stored at 20°C (blue line) and 5°C chilled and rewarmed to 20°C (red line). The 12.5°C (green dots) and 5°C (yellow dots) were also shown, but due to the low production they are not fitted into a curve.

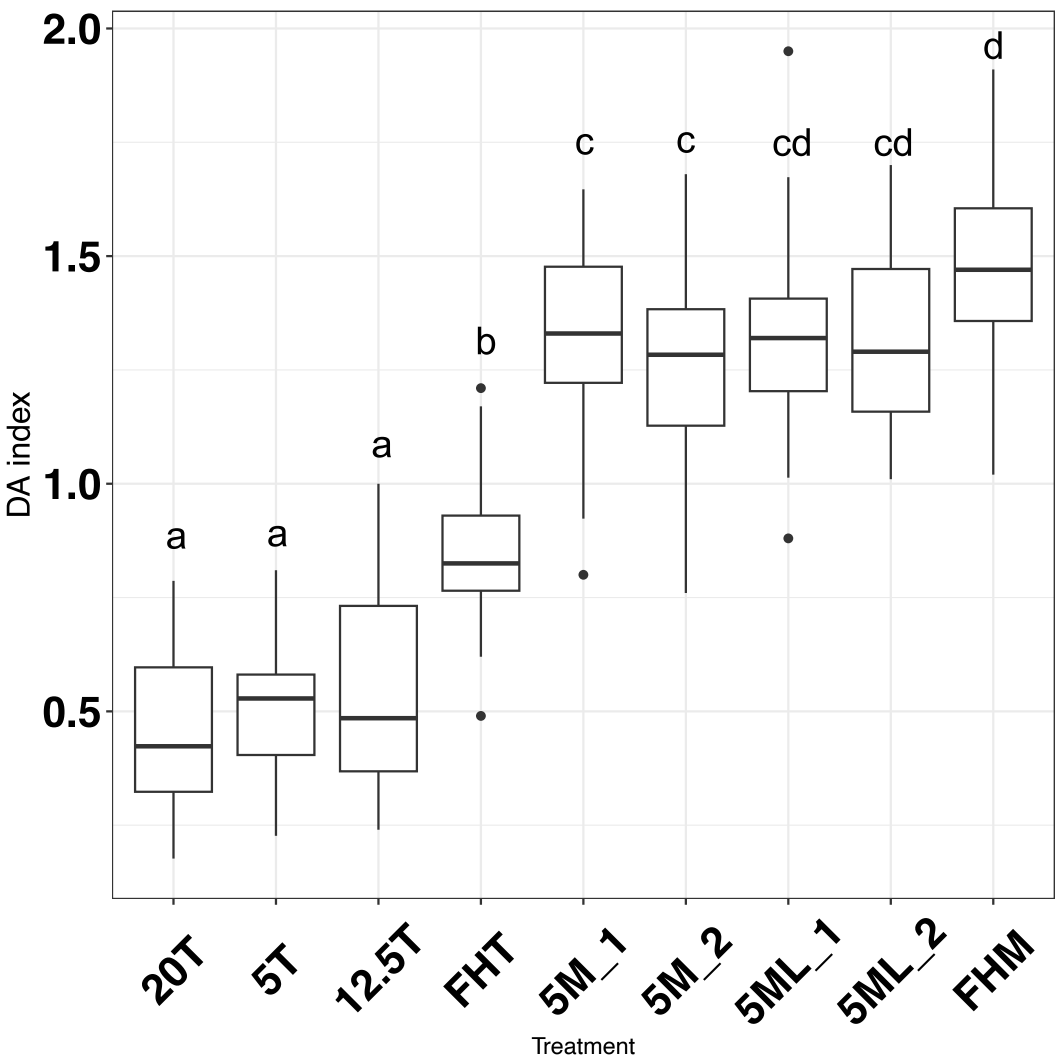


Figure S12 Boxplot of the postharvest fruit DA index. ‘5M’- mature green fruit were chilled for two weeks in the dark condition (same as all other analysis in this work), and the ‘5ML’ were supplemented with the normal white light. The DA measurement of the ‘5M’ and ‘5ML’ were repeated shown as ‘5M_1’, ‘5M_2’, ‘5ML_1’ and ‘5ML_2’.

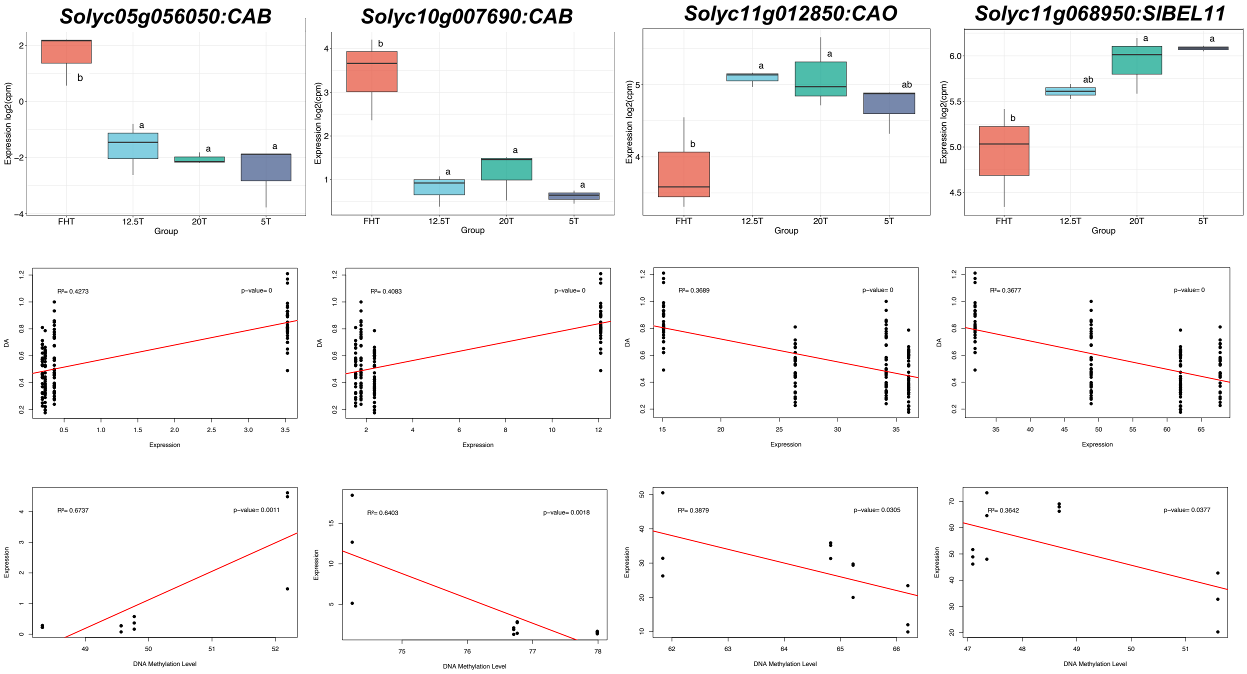


Figure S13 Photosynthetic genes with significant correlation and difference in expression. The four columns are for the individual gene's expression box plot, correlations for DA index and gene expression, and correlations for gene expression and DNA methylation levels.

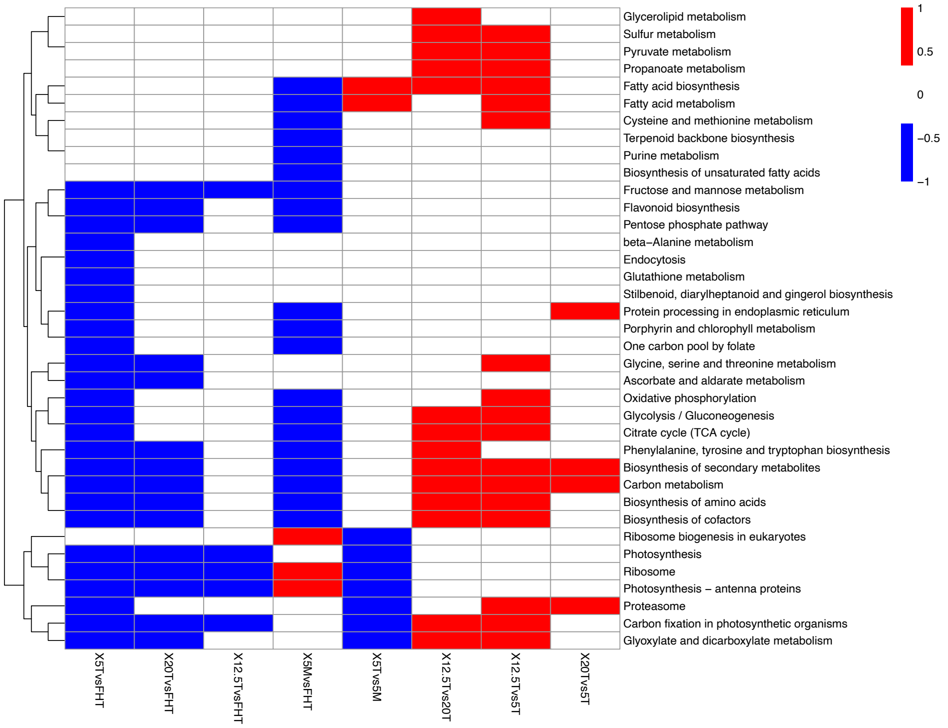


Figure S14 Transcriptomic analysis by KEGG annotation. The boxes in red, blue or white represent upregulation, downregulation, or not significant pathways (FDR< 0.01).

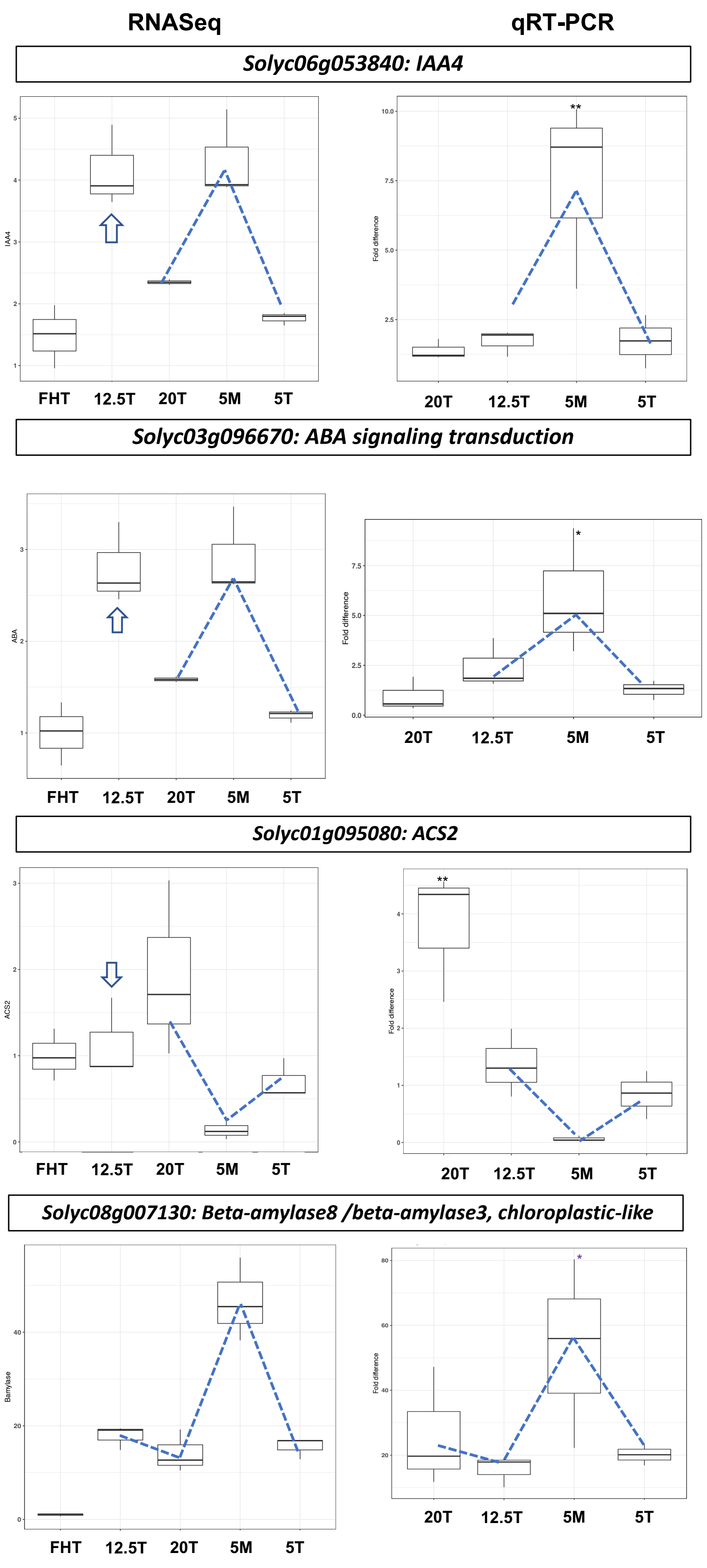
RNASeq
qRT-PCR
Solyc06g053840: IAA4
IAA4
Fold difference
FHT
12.5T
20T
5M
5T
**
Solyc03g096670: ABA signaling transduction
ABA
Fold difference
*
Solyc01g095080: ACS2
ACS2
Fold difference
**
Solyc08g007130: Beta-amylase8 /beta-amylase3, chloroplastic-like
Bamylase
Fold difference
*

Figure S15 qRT-PCR validation of the selected DEGs from RNASeq. Single asterisk (*) and double asterisks (**) refer to significant differences between 'FHT' and the postharvest treatment with *p-value* < 0.05 and *p-value* < 0.01, FHT were set as 1. The blue dashed lines indicate the trend of gene expression across samples. Compared to the 'FHT', the '20T', '5M' and '5T' groups are highly consistent between the two analysis methods, while the '12.5T' is differentially expressed for some genes shown as the blue arrows.
